# R-loop editing by DNA cytosine deaminase APOBEC3B determines the activity of estrogen receptor enhancers

**DOI:** 10.1101/2022.10.21.513235

**Authors:** Chi Zhang, Yu-jing Lu, Bingjie Chen, Zhiyan Bai, Alexia Hervieu, Marco P. Licciardello, Mei Wang, Costas Mitsopoulos, Bissan Al-Lazikani, Marcello Totorici, Olivia W. Rossanese, Paul Workman, Paul A. Clarke

**Affiliations:** Centre for Cancer Drug Discovery, the Institute of Cancer Research, London, United Kingdom; Shanghai Institute of Biological Products, Shanghai, China; Institute of Biomedical and Pharmaceutical Sciences, Guangdong University of Technology, Guangzhou, China; Agricultural Genomics Institute at Shenzhen, Chinese Academy of Agricultural Sciences, Shenzhen, China; GMU-GIBH Joint School of Life Sciences, Guangzhou Medical University, Guangzhou, Guangdong, China

**Keywords:** APOBEC3B, estrogen receptor, R-loop, TDRD3, DNA deamination, double-strand break, DNA damage repair

## Abstract

Estrogen receptor (ER) activation results in the formation of DNA double strand breaks (DSB), which promote genomic instability and tumour heterogeneity in ER-positive breast cancers. The single-stranded DNA (ssDNA) cytosine deaminase APOBEC3B (A3B) regulates ER activity by inducing DSB at ER enhancers. To delineate how A3B recognises its substrates and unveil the underlying mechanism leading to the formation of ER-induced DSB, we sampled A3B-mediated deamination sites using whole genome sequencing in a human breast cancer cell model lacking base excision repair function. Our genome-wide analysis revealed that C>U conversions carried out by A3B in R-loop structures are processed into DSB in the vicinity of ER promoters or enhancers. A mechanism which required both the processing of A3B-editing sites and R-loops by distinct DNA damage repair mechanisms. In addition, using BioID-enabled mass-spectroscopy proteomics, we identified TDRD3 as a key A3B-binding partner directing the activity of A3B to ER-induced R-loops. This study suggests a function for A3B in sustaining tumour evolution as an adaptive response at the transcriptional and epigenetic level and supports A3B as a promising target to control ER activity in cancer.

## Introduction

Estrogen (17-β E2; E2) is a hormone that is essential for the development and maintenance of the human reproductive system and secondary sexual characteristics. Prolonged exposure to E2 contributes to the development of breast, ovarian and uterine cancers^1^. E2 binds and activates the two estrogen receptor (ER) isoforms, ERα (*ESR1*) and β (*ESR2*) which are expressed in different tissues, but can form both homo- and heterodimers when co-expressed^2^. Upon E2 binding, ER translocates to the nucleus and acts as a key transcription factor by recognising E2 responsive elements (EREs) in genomic DNA. Translocation of ER to the nucleus and binding of ER to EREs leads to altered gene expression of thousands of downstream genes^3^. The gene set regulated by ER activation encompasses genes encoding essential reproductive functions, including cell proliferation and survival, and is linked to cancer development when ER is inappropriately activated^4^. In addition to this role as a regulator of gene expression, ER can also promote the development of cancer by inducing DNA damage^5^. In ER-expressing breast epithelial cells, short-term stimulation by E2 triggers the formation of DNA double-strand breaks (DSBs)^6, 7^, which contributes to chromosome instability and aneuploidy^8^. Increased genomic damage is particularly apparent in ER-positive breast cancers, where there is a high degree of chromosome instability characterized by mutations, gene copy number alterations and recombinations^9, 10^. It has been hypothesised that ER-induced DNA damage is pivotal to the activity of ER since the DNA repair pathway elements not only participate and promote transcription activation, as exemplified by transcription-coupled nucleotide excision repair (TC-NER) factors^11, 12^, but also foster favourable conditions for transcription^13^ such as DNA demethylation^14^, chromatin remodelling ^15^ and enhancer-associated RNA (eRNA) synthesis^16^.

To date, several models have been proposed to account for the generation of E2-induced DSBs, including: the action of DNA topoisomerase IIβ^17^, formation and processing of co-transcriptional R-loops^7^, and DNA damage triggered by deamination editing^6^. In recent years there has been particular interest in the role of apolipoprotein B mRNA-editing enzyme catalytic polypeptide-like 3 (APOBEC3B, A3B) and its role in multiple human cancers^18^. A3B is a member of the APOBEC3 enzyme family, which comprises seven closely related DNA deaminases that catalyse cytosine-to-uracil (C>U) editing of single-stranded DNA (ssDNA)^19^ or RNA. In addition to their normal functions as immune defences against DNA viruses and transposons ^20, 21^, APOBEC3 catalytic activity has been reported to drive genetic heterogeneity in breast cancer by inducing C>T transitions and C>G transversions at 5’TCW motifs (W = A or T)^9, 22, 23^. Although mounting evidence is pointing APOBEC3A, a highly homologous enzyme to C-terminal A3B, as the main contributor to APOBEC-driven mutation burden and cancer evolution^24^, A3B-driven cytidine deamination still has a prominent role in cancer due to two facts. First, as the only nuclear-localised member of the APOBEC3 family, elevated A3B expression is reported in >50% of primary breast cancers^22, 23^; and secondly, A3B binds directly to ERα and edits ER binding sites, which triggers DNA damage response (DDR) and are then processed by DNA base excision repair (BER) and non-homologous end joining (NHEJ) into DSBs^6^. The right amount of DSBs, together with the subsequent epigenetic modifications, determines the transactivation activity of ER^6^. Loss of functional A3B ablates ER-driven cell proliferation and prolongs the therapeutic benefit of tamoxifen in ER-positive xenograft models^25^. These data support A3B as a regulator of ER, and also as a promising target to tackle drug resistance in ER-targeting therapies.

Elevated transcriptional activity leads to the formation of chromatin-associated R-loop structures, which are formed by re-annealing of nascent RNA with the transcribed strand of DNA and results in a three-stranded structure consisting of a DNA:RNA hybrid and a displaced ssDNA strand^26^. Previous work has shown that clearance of co-transcriptionally formed R-loops by DDR machinery is a main source of DSBs caused by ER activation^7^. Since the displaced ssDNA in R-loops could serve as a substrate for A3B, it has been proposed that A3B could be involved in R-loop-associated functions. This hypothesis is supported by the use of APOBEC-Cas9 engineered system as tools to edit R-loop DNA cytosines^27^, and the fact that activation-induced cytidine deaminase (AID), a member of the APOBEC/AID deaminases family, edits R-loops to facilitate antibody gene hypermutation and class switch recombination in B lymphocytes^28^. More recently, A3B has been found to influence R-loop homeostasis^29^. However, despite these endeavours, it is unknown whether R-loop editing by A3B is directly associated with its function as an ER coactivator.

An ideal way to address the potential role of R-loop editing by A3B on the co-activation of ER-regulated transcription is to sample A3B editing sites on a large scale using ER-dependent models. This approach would reveal potential hot spots of A3B editing, which could be used to infer the mechanism that governs A3B-dependent DNA editing and point to how A3B regulates ER activity. Large-scale whole-genome sequencing (WGS) studies performed on cell lines and primary samples have identified sites of APOBEC editing^10, 30^. However, these studies may be confounded by a ‘survivorship bias’ where the detected A3B-edited sites are limited to those that were not repaired by DDR, as it is known that BER removes U:G mispairing resulting from A3B editing^24, 31^. This caveat may also affect studies employing A3B-overexpression cell models^32–35^. To circumvent this, we sampled A3B editing sites using an engineered BER-deficient breast cancer cellular model and used a multi-omic approach to elucidate the role of A3B in the molecular mechanism of ER-regulated gene expression.

## Results

### Detection of A3B deamination sites using a BER-deficient human breast cancer cell model

In order to capture A3B deamination sites that may have a functional impact on protein coding or regulatory DNA sequences, we performed whole-genome sequencing (WGS) of ER-positive T-47D human breast cancer cells co-expressing doxycycline-inducible A3B. and humanised bacteriophage PBS2 uracil glycosylase inhibitor (hUGI) peptide (**Figure 1A**, top panel) to inhibit the action of uracil-DNA glycosylase (UDG/UNG) and prevent excision of the A3B-catalysed uracil bases^22, 24^. The T-47D cell line was chosen because it does not express APOBEC3A which may confound the study^6^, and also carries a loss-of-function *TP53* mutation that is known to protect the cells from the synthetic lethality caused by ectopic expression of A3B^36^. After confirming the inducible expression of A3B/hUGI by immunoblotting, we exposed the cells to doxycycline for five days (**Figure S1A-B**), and sequenced genomic DNA samples from five individual T-47D clones (A+ colonies). As controls, sequencing results from three un-induced colonies (A- colonies) and three T-47D colonies (M+ colonies) co-expressing hUGI and enzymatically-dead A3B E68Q/E255Q double mutant (A3B**)^6^ were used to represent single-nucleotide variants (SNV) arising either from the clonal separation process or base-editing by endogenous ABOBEC/AID-family enzymes (**Figure 1B**). Joint mutation calling by the GATK4 computational programme was carried out using a parental T-47D reference sample, the result of which showed a significantly elevated proportion of non-CpG C>T mutations in the A+ colonies over the M+ and A- colonies (**Figure 1C**). We next assessed the extent of focal hyper-mutation among the detected mutations using both intermutational distance^9^ (**Figure 1D, S1C**) and an algorithm that detects somatic local hyper-mutation (HyperClust^37^) (**Figure 1E**). Unlike previous studies, where kataegis-like mutation clusters were prevalent in BER-functional models^32, 34^, fewer clustered mutational events were detected in A+ colonies where BER activities were suppressed by hUGI. However, A+ colonies contained more non-recurrent and diffused form of clustered mutation termed omikli^37^ (**Figure 1E**). Analysis of the resulting SNVs against pre-defined COSMIC single-base substitution (SBS) signatures (version 2) using non-negative matrix factorization (NMF)^30^ identified the enrichment of signature SBS2 which was attributed to the activities of AID/APOBEC family of cytidine deaminases (**Figure 1F**). *De novo* NMF signature extraction revealed two prevalent signatures among the sequenced control colonies: signature A which is more prevalent in A-/M+ colonies and might account for the variation among T-47D subclones, and signature B which is mostly enriched in A+ colonies and harbours the A3B-preferred 3’-TCW trinucleotide motif, resembling the mutational process attributed to A3B editing (**Figure 1G, H**). Notably, other than C>T substitutions, a very weak contribution by C>A and C>G substitutions at the 5’-TC motif were observed in signature B, suggesting that most of the A3B-mediated uridine sites were subjected to BER processing. In addition, both SBS2 and signature B are also found in M+ colonies, but enriched to a much lesser extent, implying the capture of intrinsic APOBEC/AID-family-mediated mutations by BER blockade. These data demonstrated that the BER-deficient breast cancer model successfully preserves and captures A3B activity on genomic DNA in the absence of the interference of downstream DNA repair mechanisms.

**Figure 1:**
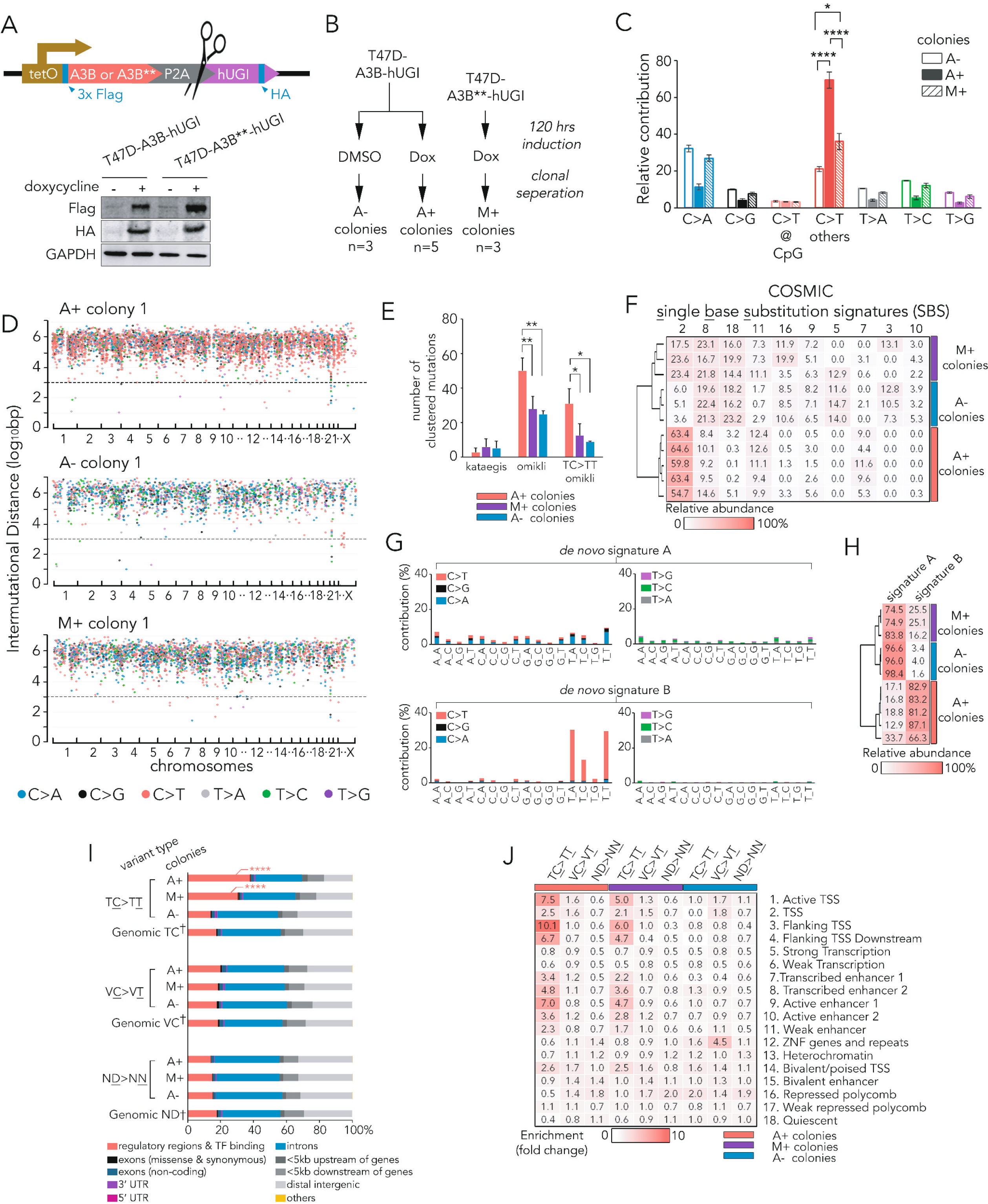
Detection of A3B deamination sites in BER-deficient cell models. (A). Lentiviral inducible system used in this study and immunoblotting of cells with and without 24-hour induction by 1 μg/ml doxycycline. A3B** denotes A3B with E68Q/E225Q mutations. (B). Schematic of sample preparations for WGS studies. (C). Mean relative contribution of the indicated types of point mutations for the three indicated sample groups. Error bars = SD; *: p < 0.05; ****: p < 10^−4^, one-way ANOVA. (D). Representative waterfall plot of intermutational distance (IMD) of each mutation identified in the three indicated groups of colonies. Dotted line denotes IMD ≤ 10^3^ bp. (E). Mutation clusters detected by HyperClust for three sample groups. Data for mean value and error bars for SD. *: p < 0.05; **: p < 0.01, one-way ANOVA. (F). Heat map showing the cosine similarity scores of top-ranking COSMIC signatures for each indicated samples. (G). Relative contribution of each indicated trinucleotide alternation to the two top-ranking *de novo* mutational signatures identified by NMF analysis. (H). Heat map showing the cosine similarity scores of mutational signatures in G for each indicated samples. (I). Distribution of variant consequence as determined by VEP for indicated types of variants. †: random sampling of 10^6^ sites. ****: p < 10^−4^, χ^2^ test comparing to random sites. (J). Heat map showing enrichment score of chromatin states for ± 50 bp regions flanking point mutations. Chromatin states were derived by ChromHMM (**Figure S1D**).

We next explored the functional implications of the captured A3B-editing sites using the Ensembl Variant Effect Predictor (VEP)^38^. When compared to random genomic locations, the TC>TT substitution in A+ and M+ colonies was significantly enriched in non-coding regulatory and transcription factor (TF) binding regions relative to protein-coding regions, an effect not seen in either VC>VT or ND>NN substitutions (**Figure 1I**). To further dissect function of these regulatory regions, we constructed an 18-state ChromHMM model for the T-47D cells using ChIP-seq data from six histone marks (**Figure S1D**) and computed its degree of overlap between the captured mutations (**Figure 1J**) using previously described methods^39^. In the A+ colonies we found overrepresentation of TC>TT mutations at transcription start sites (TSS) and enhancers, especially at the flanking regions of TSS and active enhancers. This pattern can also be found for intrinsic APOBEC-induced mutations in M+ colonies, but to a lesser extent than the A+ colonies. Analysis of DNA replication timing analysis using T-47D Repli-seq data from ENCODE indicated that the mapped TC>TT substitutions and omikli events in A+ colonies occurred early in DNA replication (**Figure S1E**). These findings are in accordance with previous observations that A3B targets genomic regions being actively transcribed and/or involved in early replication^37^. Overall, our data demonstrate a clear association between A3B-mediated base-editing and sites of gene transcription, consistent with a previously defined role for A3B in transcription regulation^6^.

### A3B binds and deaminates ssDNA in R-loops

We evaluated whether R-loops, a co-transcriptionally formed DNA:RNA hybrid, could provide substrates for A3B activity. In A+ colonies, TC>TT substitutions are flanked by a high level of GC-skewness for DNA sequences, which denotes overrepresentation of guanine bases in the same DNA strand as the TC>TT editing event (**Figure 2A**). This also translates to an enrichment of G-quadruplex-forming sequences in ±500 bp regions around A3B-editing sites (**Figure 2A**). A similar effect can be seen around TC>TT substitutions from M+ colonies, albeit to a much lesser extent, but not around TC>TT or other di-nucleotide substitutions in the A- colonies (**Figure 2A, S2A-B**). Both GC-skew and G-quadraplex forming sequences are features reported to facilitate the formation or stabilisation of R-loops^40^. We mapped R-loops in parent T-47D cells using strand-specific DNA:RNA immunoprecipitation sequencing (ssDRIP-seq; **Figure S2C**) coupled with RNase H-dependency, GC-skewness and MNase-seq analyses to verify the identification of R-loops (**Figure S2D-E**). We found significant enrichment of R-loop signals in close proximity to TC>TT substitutions in A+ colonies. Strand-specific sequencing demonstrated that the TT>TC substitutions occurred on the ssDNA strand of the R-loop consistent with A3B preferentially editing the displaced ssDNA strand (**Figure 2B**). A weaker R-loop signal can be found at TC>TT substitutions from M+ colonies, but not in any other types of di-nucleotide substitutions (**Figure S2F**).

**Figure 2:**
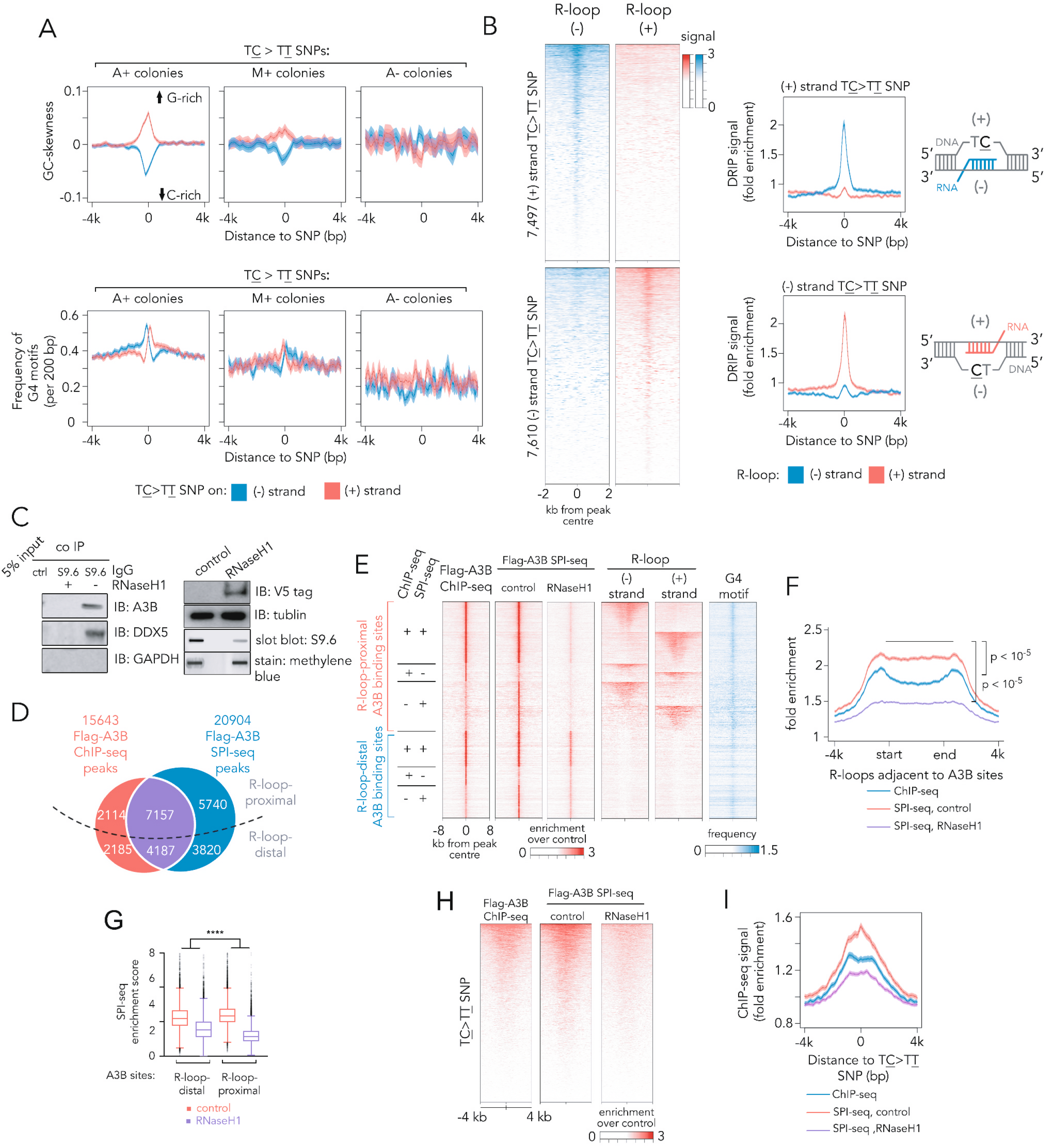
A3B binds and edits R-loop. (A). Profiles of GC skewness and frequency of G-quadruplex (G4) motifs in regions flanking the TC>TT SNP. Line denotes average value and shaded areas 95% CI. (B). Heat maps showing ssDRIP-seq signals in regions flanking the TC>TT SNP identified in A+ colonies (left), their corresponding signal profile plots (middle), and schematics demonstrating the position of R-loop in relation to the site of TC>TT SNPs (right). (C). S9.6 co-immunoprecipitation assay of T-47D cells transfected with or without RNaseH1-encoding vectors for 24 hours (left), and representative immunoblots and slot blots from input samples (left). (D). Venn-diagram showing the overlap of Flag-A3B binding sites from ChIP-seq and SPI-seq. Numbers of sites proximal to (within 1.5 kb) or distal to R-loop are shown. (E). Heat maps of enrichment signal from indicated sequencing experiments at the flanking regions of consensus Flag-A3B ChIP-seq and SPI-seq peaks. (F). Profile of Flag-A3B ChIP-seq or SPI-seq signal for R-loops that are proximal to A3B sites. Line denotes average value and shaded areas 95% CI. Statistic represents one-way ANOVA using enrichment scores at indicated regions. (G). Tukey boxplots showing enrichment scores from SPI-seq experiments. ****: p ≤ 10^−4^, two-way ANOVA assessing the effect of R-loop proximity on response to 24-hour RNaseH1 overexpression. (H). Heat maps showing Flag-A3B ChIP-seq or SPI-seq signal in regions flanking TC>TT SNP identified in A+ colonies. (I). Signal profile for Flag-A3B ChIP-seq or SPI-seq signal in regions flanking TC>TT SNP identified in A+ colonies. Line denotes average value and shaded areas 95% CI.

To further examine the association between R-loops and A3B-mediated editing, we first carried out co-immunoprecipitation experiments with the R-loop-binding S9.6 antibody, a previously reported methodology for isolating R-loops and their binding proteins^41^. Similar to DDX5, a previously validated R-loop binding protein^41^, A3B co-complexed with immuno-precipitated R-loops, which was abrogated by the overexpression of RNaseH1, an enzyme that digests RNA:DNA hybrids (**Figure 2C**). Next, the genomic binding sites of flag-tagged A3B were mapped using chromatin immunoprecipitation sequencing (ChIP-seq) and ssDNA-associated protein immunoprecipitation (SPI-seq)^42^ the latter of which is an approach better suited to capturing protein-bound ssDNA. More than half of A3B peaks detected by either method are found in the vicinity (< 1.5 kb) of R-loops. Although both methods yielded a similar proportion of R-loop-proximal A3B peaks (61% of all sequences with SPI-seq, 59% with standard ChIP-seq) the SPI-seq approach was more sensitive (p < 0.01, χ^2^ test, **Figure 2D-E**). We also found that the SPI-seq captured more A3B-bound ssDNA fragments located in the body of R-loops, in contrast to standard ChIP-seq which captures R-loop-flanking dsDNA (**Figure 2F**). Induced expression of RNaseH1 is known to reduce genomic R-loop content^7^, and we found that A3B peaks adjacent to R-loops were specifically reduced by expressing RNaseH1 (**Figure 2E-G**), demonstrating a negative effect of R-loop clearance on A3B binding to genomic DNA. In addition, we observed a higher level of SPI-seq signal at TC>TT substitutions in A+ and M+ colonies compared with conventional ChIP-seq, and the binding of A3B to these sites was modulated by the expression of RNaseH1 (**Figure 2H-I, S2G**). This suggests R-loop formation facilitates A3B binding and cytidine deamination.

### ER-induced R-loop formation facilitates A3B binding

Since activation of ER triggers the formation of co-transcriptional R-loops^7^, we hypothesised that elevated levels of genomic R-loops would boost the availability of A3B substrate and in turn facilitate A3B binding to ER-activating enhancer regions. To test this, flag-tagged A3B binding sites and R-loops that are responsive to E2 stimulation were mapped using SPI-seq and ssDRIP-seq (**Figure 3A**) in T-47D breast cancer cells, which allowed the study of their spatial-temporal relationship. Similar to previous reports^6, 7^, we observed an overall increase in A3B binding (23,711; 53% of all the peaks) and R-loop (18,567; 16% of all the peaks) peaks upon exposure to E2 for two hours (**Figure 3B**). Notably, following exposure to E2, a significant proportion of newly formed A3B binding sites are proximal to R-loops (p < 10^−5^, χ^2^ test), and correspondingly, R-loops near A3B binding sites are more susceptible to E2 induction (**Figure 3B**).

**Figure 3:**
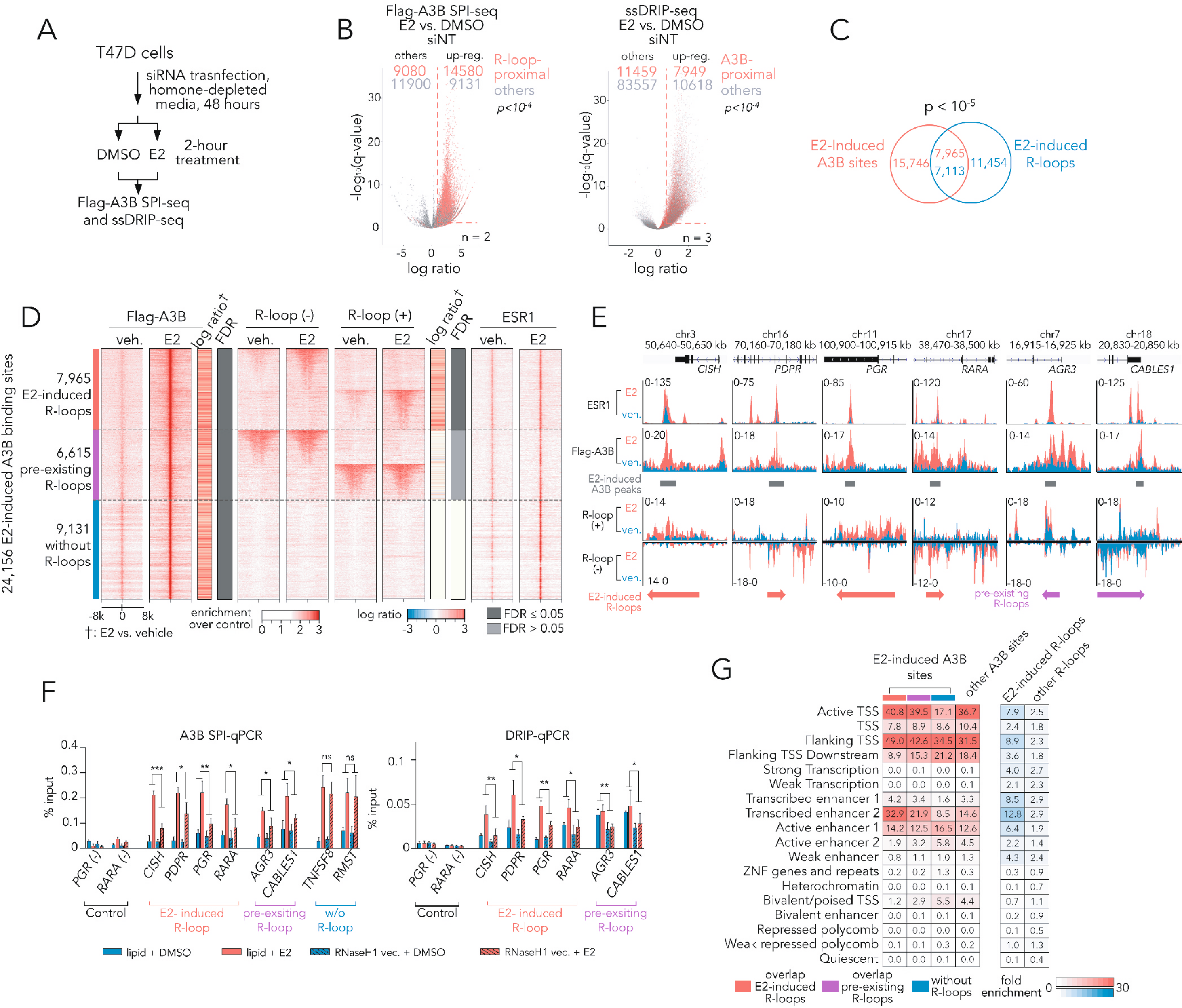
R-loops induced by ER activation facilitates A3B binding. (A). Schematic of the quantitative A3B SPI-seq and experiment carried in this study. (B). Volcano plot summarising the effect of two-hour E2 stimulation on Flag-A3B binding sites determined by SPI-seq (left) and R-loop sites determined by ssDRIP-seq (right). R-loop-proximal Flag-A3B sites are coloured in red. Up-regulated sites are defined as FDR ≤ 0.05 with fold change ≥ 1.5 (indicated by red dotted line). P value represent statistical significance by χ^2^ test of non-random association between R-loop or A3B proximity and response to E2. (C). Venn diagram showing the overlap between E2-induced Flag-A3B binding sites and E2-induced R-loops. P value was obtained by χ^2^ test for the independence of the two sets in comparison with the genomic background. (D). Heat maps showing Flag-A3B SPI-seq, ssDRIP-seq and ESR1 ChIP-seq signals in ±8 kb regions flanking E2-induced Flag-A3B binding sites. (E). Representative signal tracks for ESR1 ChIP-seq, Flag-A3B SPI-seq and ssDRIP-seq at E2-induced Flag-A3B binding sites that are proximal to E2-induced R-loops (CISH, PDPR, PGR and RARA) or R-loops pre-existing to E2-induction (AGR3 and CABLES1). (F). Bar graph showing Flag-A3B SPI-qPCR and DRIP-qPCR results. Data represent mean of three biological replicates and error bars for SD. *: p ≤ 0.05; **: p ≤ 0.01; ***: p ≤ 0.001; ns: p > 0.05, two-way ANOVA assessing the effect size of RNaseH1 treatment on E2 response. (G). Heat map showing enrichment scores for ChromHMM chromatin states at indicated A3B binding sites or R-loops.

Because of the significant overlap between E2-induced A3B binding sites and R-loops (**Figure 3C**), we investigated a potential causal relationship between enhanced A3B binding and formation of R-loops. Depletion of A3B with siRNA does not affect the overall levels of R-loop signal or the increase in R-loops in response to E2-induction, as detected using slot blotting with S9.6 antibody (**Figure S3A-C**). Quantitative ssDRIP-seq also shows an increase of R-loops in proximity to A3B binding sites despite knockdown of A3B (**Figure S3D**). Using DESeq2 quantification method^43^, we investigated the interaction between A3B siRNA and E2 treatments and observed minimal combinatorial effect on R-loop up-regulation between the two treatments (**Figure S3E**). More than half of E2-induced A3B peaks were situated either near E2-induced R-loops (34%) or R-loops that exist prior to E2 treatment (28%, **Figure 3D**), while R-loop-proximal A3B peaks had higher degree of induction upon E2 treatment relative to R-loop-distal A3B peaks (**Figure S3F**). These results indicate that elevated A3B binding is unlikely to be linked to induction of R-loops and instead that R-loop availability may determine E2-induced A3B binding and editing. To test this, we investigated whether overexpression of RNaseH1 affects A3B binding in five loci with well-defined ER binding and R-loop formation (**Figure 3E**). In contrast to A3B binding sites without R-loops, clearance of R-loops upon overexpression of RNaseH1 impairs A3B binding induced by exposure to E2 (**Figure 3F**). These data confirm that the binding of A3B to specific regions induced by exposure to E2 involves a subset of R-loops serving as substrates.

Our findings suggest a model where R-loop formation reshapes the genomic landscape around EREs to facilitate recruitment of A3B. Using our ChromHMM model, we found that E2-induced A3B binding sites that overlap with R-loops have higher enrichment scores in TSS, TSS flanking regions, and transcribed enhancers relative to those E2-induced A3B binding sites that are distal to R-loops (**Figure 3G**). Similarly, R-loops induced by E2 exposure are enriched in TSS and transcribed enhancers compared to the remaining pool of the non-induced R-loops, suggesting co-transcriptionally formed R-loops by E2 induction may facilitate A3B binding. Analysis of TF binding motifs reveals a different set of TFs potentially enriched in E2-regulated R-loop proximal peaks as opposed to R-loop-distal A3B peaks, where most sequence motifs in the former group are G-rich (**Figure S3G**). Collectively, our data suggest that enhanced R-loop availability induced by transactivation at ER enhancers or promoters may regulate A3B binding and editing activity.

### A3B promotes DSB formation at E2-induced R-loop

Previous studies have shown that DSB formation induced by ER transactivation can be arise from the processing of either co-transcriptionally formed R-loops or A3B editing sites by DNA damage repair mechanisms^6, 7^. To investigate the link between these processes, we employed DSBCapture-seq to precisely map genomic DSB sites^44^, and to study the dependency of E2-induced DSBs on A3B and/or R-loops. We exposed T-47D breast cancer control and A3B-depleted cells to either DMSO or E2 for two hours, followed by DSBCapture-seq procedures (**Figure 4A**). Differential peak analysis of the sequencing results by DiffBind^45^ revealed two groups of DSBs: DSBs that are induced by exposure to E2 (‘E2-induced DSBs’), and DSBs modified by A3B knockdown (‘A3B-modified DSBs’, **Figure 4A**). Our results echo the reported widespread DNA damage following by ER activation^5–7, 46^, and show that the 15,259 E2-induced DSB peaks accounts for 30% of all the detectable DSBs in the genome of the T-47D cell-line. Notably, 5,308 of the DSB peaks were modified by A3B-depletion, of which a majority (88%, p < 10^5^ by Χ^2^ test) overlapped with E2-induced DSBs (**Figure 4B**). This enrichment of A3B-modified DSBs in the E2-induced DSB set is consistent with previous reports^6^. Closer inspection of DSB sites identified in our study found that almost all (98.6%) of the A3B-modified DSB sites we detected overlapped with the E2-induced A3B binding sites identified by quantitative SPI-seq (**Figure 4C**), supporting a role for A3B in these DSBs. However, A3B binding alone is not sufficient to predict A3B-dependent formation of DSBs since 74% of the E2-induced A3B-independent DSBs, defined by up-regulated DSBs despite the depletion of A3B, also overlapped with A3B binding sites defined by SPI-seq. Instead, we found a requirement for R-loop formation prior to subsequent A3B binding to generate A3B-modified DSBs, since we detected a significant overlap with R-loops for these DSBs compared with E2-induced DSBs that were A3B-independent (**Figure 4C**). Accordingly, both R-loop formation and A3B binding distinguished A3B-modified DSBs from A3B-independent DSBs (96% vs 42%, **Figure 4D**) and were good indicators of the impact of A3B depletion on DSB formation (**Figure 4E**). Therefore, our results support the involvement of R-loop formation in the generation of A3B-induced DSBs.

**Figure 4:**
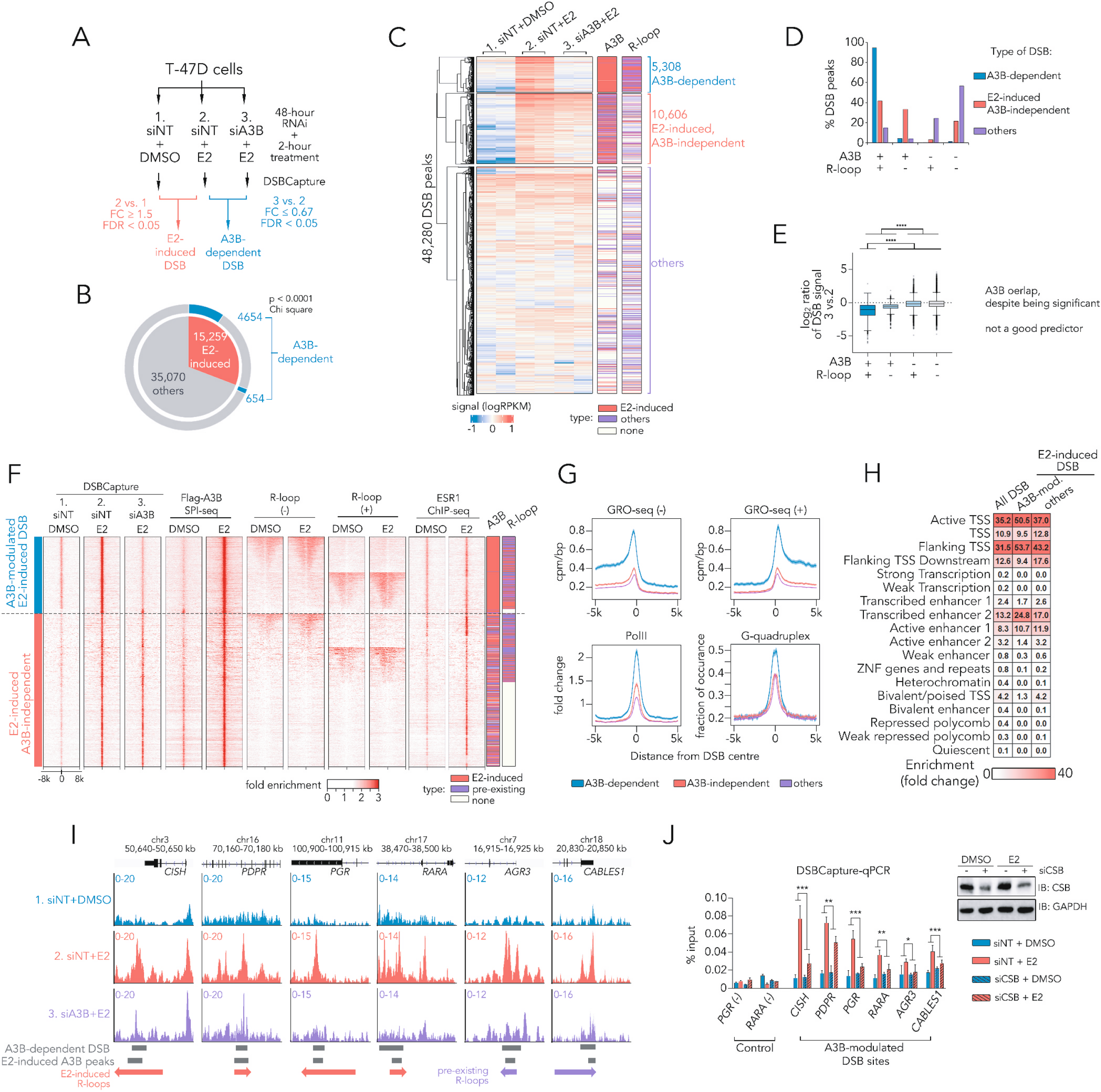
A3B promotes E2-induced DSB formation at R-loops. (A). Schematic of quantitative DSBCapture experiments to identify E2-induced and A3B-dependent DSBs in T-47D cells. (B). Pie chart showing results of quantitative DSBCapture experiments, p value derived from χ^2^ test evaluating the statistical significance of non-random association between A3B-dependent and E2-induced DSBs. (C). Heat maps visualising normalised signal for identified DSBs in T-47D cells. Each row represents one DSB site and each column a unique sample, and types of overlapping A3B or R-loop features are displayed alongside. (D). Bar graph showing percentage of DSB peaks overlapping A3B and/or R-loop. (E). Tukey boxplots showing log_2_ ratio of DSBCapture signals between A3B and non-targeted siRNA-transfected cells after 2 hours of E2 transfection. Statistical analysis denotes two-way ANOVA, ****: p ≤ 10^−4^. (F). Heat maps for DSBCapture, A3B SPI-seq, ssDRIP-seq and ESR1 ChIP-seq signals flanking ±8 kb of centre of DSBs. Also shown are bars indicating types of overlapping Flag-A3B SPI-seq and ssDRIP-seq peaks. (G). Profiles of GRO-seq, PolII ChIP-seq signal and G-quadruplex occurrences in regions flanking indicated types of DSBs. Line represents average signal or frequency and shaded area denote 95% C.I. (H). Heat map showing enrichment scores for ChromHMM chromatin states at indicated DSBs. (I). Representative signal tracks for DSBCapture at E2-induced Flag-A3B binding sites that are proximal to E2-induced R-loops. (J). Bar graph showing DSBCapture-qPCR results. T-47D cells were treated with NT or CSB siRNA for 48 hours before 2-hour stimulation by DMSO or 100 nM E2. Data represent mean of three biological replicates and error bars for SD. *: p ≤ 0.05; **: p ≤ 0.01; ***: p ≤ 0.001; two-way ANOVA assessing the effect size of CSB siRNA on E2 response.

A closer inspection of the regions flanking DSBs revealed some distinct features associated with A3B-dependent DSBs. These included higher bidirectional transcription activities (as determined by GRO-seq), RNA polymerase II (PolII) occupancy, and denser G-quadruplex-forming sequences, all of which favour the formation of R-loops^47^ (**Figure 4F**). Using our ChromHMM model, A3B-dependent DSBs were found to be more enriched in active/flanking TSS and transcribed enhancers, a group of chromatin states also enriched in E2-induced R-loops, suggesting an influence of R-loops on the DSB landscape similar to that seen for A3B binding sites (**Figure 4G**). These observations indicated a dependency on R-loops during the processing of A3B editing sites to DSBs. To test this, we analysed the five R-loop-overlapping A3B binding sites where A3B-dependent DSBs were found (**Figure 4H**) and tested the effect of RNAi-mediated depletion of Cockayne syndrome group B (CSB) protein. CSB is a TC-NER component known to be required for converting R-loops into DNA strand breaks^48^ . Using quantitative PCR on DSBCapture-enriched DNA samples, we found that even modest CSB depletion was sufficient to reduce E2 responsive A3B-dependent DSB formation, but not E2-induced, A3B-independent DSBs (**Figure 4I**). These results revealed a requirement of R-loop processing into single-strand breaks (SSB) for the formation of A3B-dependent DSBs.

### Blocking uracil base-processing impairs E2 response

The genomic location of A3B-binding sites, especially those in proximity to R-loops and linked to the formation of DSBs, suggested a gene regulatory role for the down-stream processing of A3B base-editing. To address this, we performed RNA-seq studies on T-47D breast cancer cells harbouring a doxycycline inducible enzymatically inactive A3B** and hUGI cassette (**Figure 5A**). This construct prevents the downstream processing of the intrinsic A3B-mediated C>U editing sites by inhibiting UNG enzymes. We collected RNA samples after exposing cells to E2 for 16 hours, with or without doxycycline induction, and performed RNA-seq. For differential gene expression analysis, interaction modelling by DEseq2 programme was employed to dissect the impact of UNG inhibition (by hUGI) over E2 response^43^. Results showed that the expression of A3B**-hUGI dampened the E2-regulated response of 87% of genes (**Figure 5B**), out of which 90 genes were identified as statistically significant with an interaction model by DESeq2. (**Figure 5C**). Using pathway enrichment analysis with MSigDB hallmark gene set^49^, we observed a significant overlap between these genes and genes regulated by estrogen response pathways (**Figure 5D**), indicating a requirement for uracil repair during the transactivation of ER. In order to investigate whether the genes likely to be regulated by A3B are affected by lack of uracil-repair mechanisms, we first used the predictive algorithm rGREAT to generate two groups of genes which are most likely to associated with regions that could form gene-regulating DSBs^50^, the first one contained 86 genes that were associated with A3B-dependent DSBs, and the latter one contained 333 genes that were associated with all E2-induced A3B binding sites that were proximal to R-loops. Using our RNA-seq data as the input for leading edge analysis by gene set enrichment analysis (GSEA)^51^, we found a significant overlap between genes from both of these groups and with genes up-regulated by E2 (**Figure 5E**), suggesting the formation of A3B-dependent DSBs may facilitate the transactivation of ER-regulated genes. More importantly, the genes from both groups were also enriched with E2-regulated genes that were negatively affected by A3B**-hUGI expression which confers BER-deficiency (**Figure 5E**), suggesting that the inability to process A3B-editing sites into DSBs in certain regulatory regions hampered the transactivation of gene expression following E2 stimulation. Together, our collective results demonstrate that the downstream processing of A3B C>U editing sites, which leads to formation of DSBs, may promote E2 response in T-47D cells.

**Figure 5:**
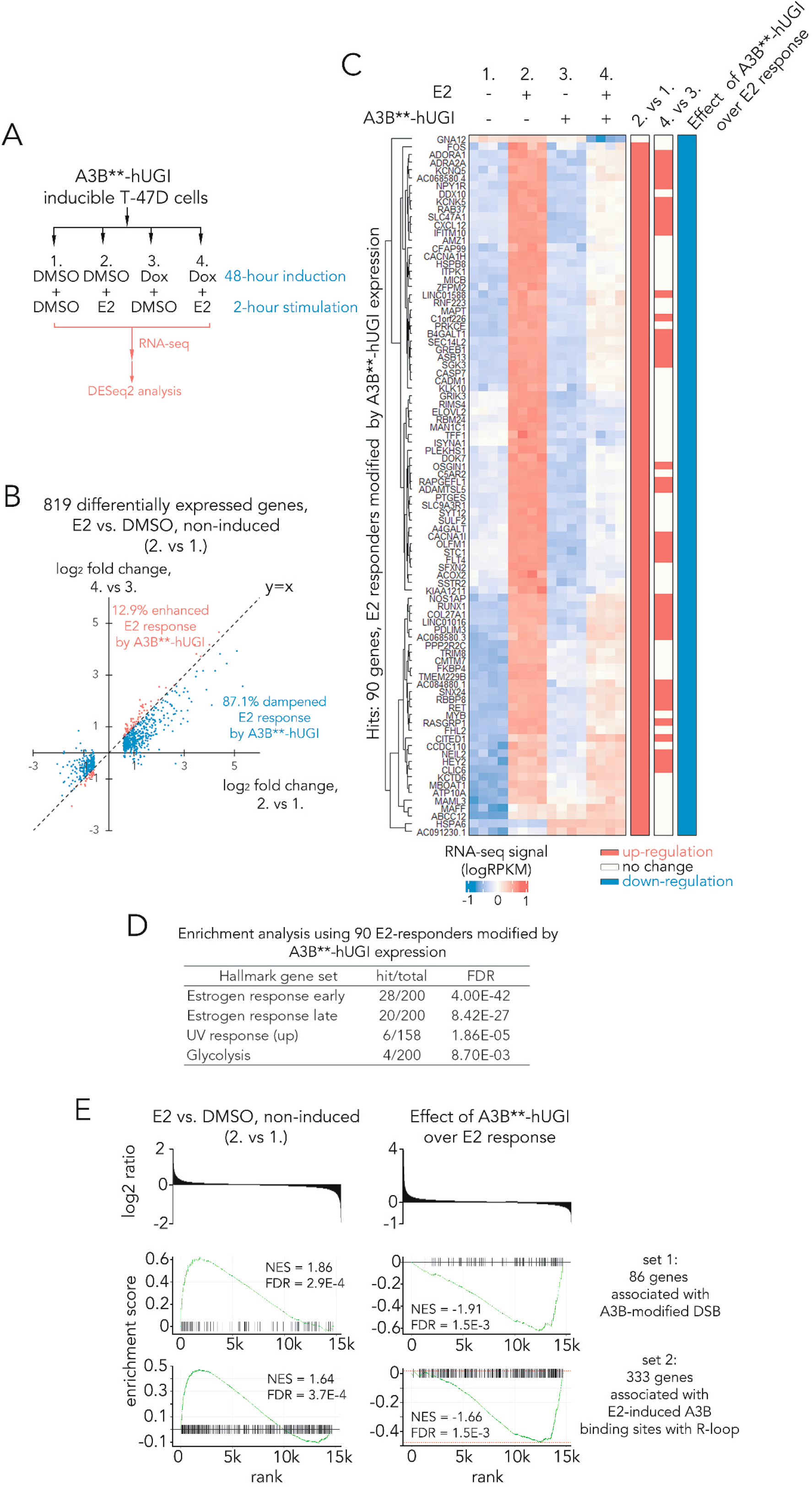
Blocking base excision repair impairs E2 response. (A). Schematic of the RNA-seq experiment in this study. (B). Dot plot showing log_2_ fold change of normalised transcript counts for 819 differentially expressed genes affected by E2 stimulation without the expression of A3B**-hUGI. (C). Heat maps showing normalised RNA-seq signals for 90 differentially expressed genes the E2 response of which were modulated by A3B**-hUGI induction. Statistical results from DESeq2 are summarised in bars on the right. (D). Table showing MSigDB hallmark gene set enrichment analysis using 90 hit genes described in (C). (E). Leading edge analysis for genes associated with A3B-dependent DSB or R-loop-overlapping E2-induced Flag-A3B binding sites. Associations of genomic regions and gene were predicted using rGREAT using a binomial Bonferroni-adjusted p value of 0.05. Leading edge analysis were conducted using fGSEA package with R.

### TDRD3 guides A3B binding to regulatory elements

We found that ER activation leads to pervasive formation of R-loops with potential to be A3B substrates; however, only a subset of these R-loops was bound by A3B, suggesting a molecular mechanism that facilitates R-loop selectivity. To test whether this potential selectivity could be driven by proteins interacting with A3B, we employed BioID^52^, a proximity-dependent labelling method, to capture A3B binding partners. A3B tagged with BirA* on either N- or C-terminal was used as bait, and BirA*-red fluorescent protein (RFP) was used as control. In addition to T-47D cells, the BioID experiments were also carried out in two additional cell lines, MDA-MB-468 with high A3B expression background, and MCF-10-A with very low A3B levels (**Figure 6A, Figure S4A**). *Bona-fide* A3B interactors were selected against common BioID backgrounds using SAINTexpress score in conjunction with historical BioID non-specific controls curated in the CRAPome database^53^. Our results showed a general overlap of A3B interactors across different cell lines, which also found a higher rate of detection in higher intrinsic A3B expressing cancer cells than the low-expression MCF-10-A cells (**Figure 6B**). Using A3B interactors detected in at least two cell lines, we plotted an interaction network using common protein-protein interaction databases (**Figure 6C**) and found two clusters of interactions: one with mainly RNA-binding or HNRNP components and the other comprising members of the TOP3B-TDRD3-FMRP (TTF) complex, a protein complex known in function in R-loop resolution within cells^54^. To validate the interactions between the TTF complex and A3B, we carried out co-immunoprecipitations and found A3B was able to interact with TDRD3 and FMRP either with or without the bridging effect of nucleic acids (**Figure 6D**). Upon closer inspection, we found that A3B binds only isoform 1 of TDRD3, but not the shorter isoforms 2 and 3, which lack part of an N-terminal OB-fold, suggesting the OB-fold site may contribute to A3B binding. TOP3B was not detected in the A3B complex, despite evidence for binding to TDRD3 at the N-terminus domain (**Fig. 6D**). As TOP3B also scored as a hit in the A3B BioID, there may be a dynamic but potentially mutually exclusive binding interaction between A3B, TOP3B and TDRD3.

**Figure 6:**
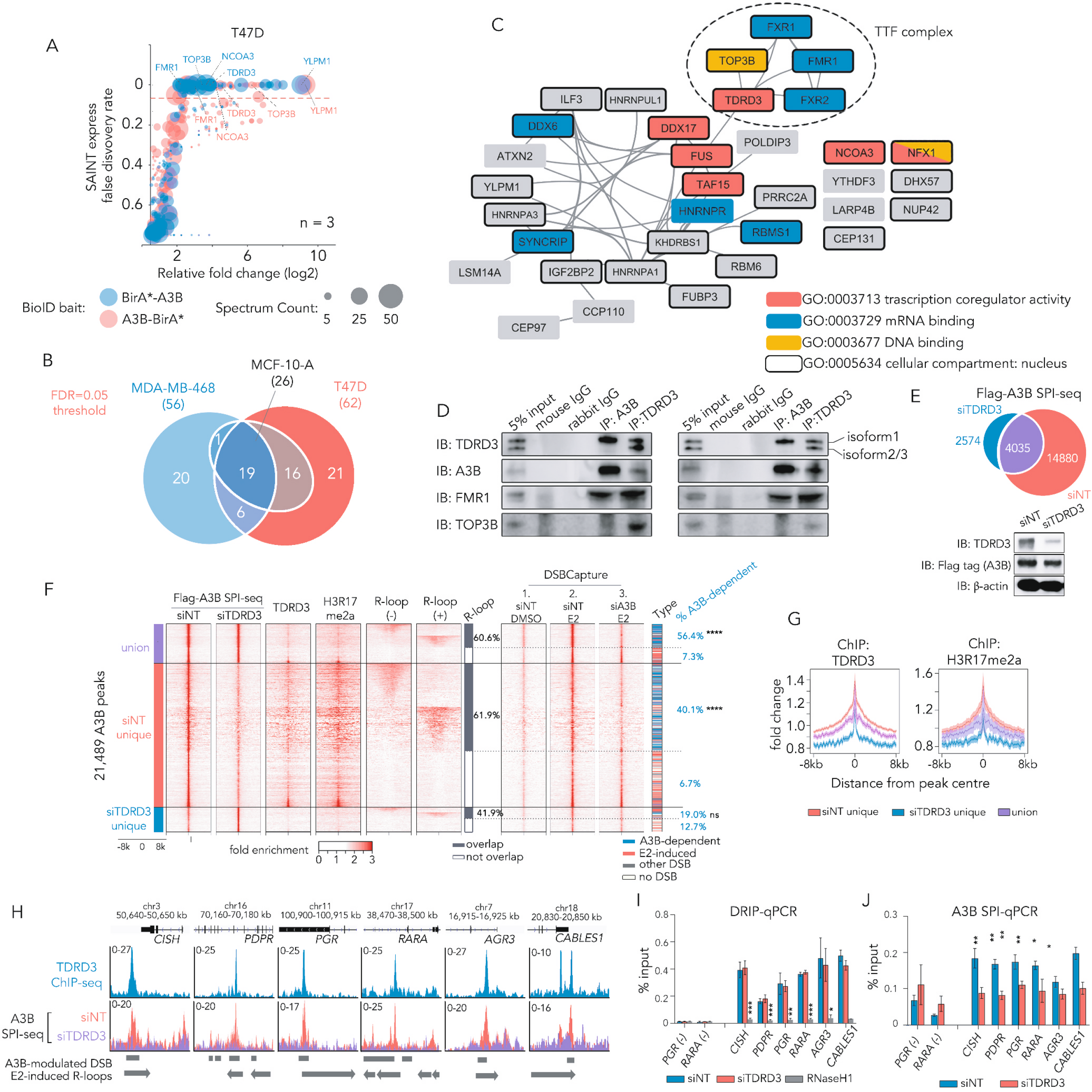
TDRD3 facilitates A3B binding to regulatory elements. (A). Bubble plot illustrating the result of BioID experiments. T-47D cells bearing lentiviral inducible BioID baits were induced with doxycycline for 24 hours before subject to BioID analysis. SAINT express FDR score were derived using BirA*-RFP pull-down results from T-47D cells as control and calculated by CRAPome. Data derived from n = 3 independent experiments and recurring hits were labelled. (B). Venn-diagram showing the overlap between BioID hits (FDR < 0.05) across three different breast cancer cell lines. (C). Network diagram showing known protein-protein interactions amongst hits that qualified in at least two different cell lines. (D). Representative immunoblotting using antibodies against A3B or TDRD3 in T-47D cells. (E). Venn-diagram showing the overlap between Flag-A3B binding sites in NT siRNA-treated or TDRD3 siRNA-treated T-47D cells, which were identified by SPI-seq. Immunoblots verifying the knockdown of TDRD3 are shown at the bottom. (F). Heat map illustrating signals from indicated sequencing experiments in ±8 kb regions flanking Flag-A3B binding sites identified by SPI-seq. Also shown are bars indicating types of overlapping DSBCapture peaks. Statistic results are from χ^2^ test evaluating the statistical significance of non-random association between overlapped R-loops and A3B-modified DSBs. **** and ns denote p ≤ 10^−4^ and p > 0.05 respectively. (G). Profiles of TDRD3, H3^R17me2a^ and CARM1 in regions flanking A3B peak centres identified by SPI-seq. (H). Representative signal tracks for TDRD3 ChIP-seq and A3B SPI-seq at indicated genomic loci. (I). Bar graphs showing results from DRIP-qPCR experiments. Samples were collected from T-47D cells transfected with NT siRNA, TDRD3 siRNA or vector encoding RNaseH1 for 48 hours. Data represent mean of n = 3 biological replicates and error bars for SD. *, ** and *** denote p < 0.05, 0.01 and 0.001 respectively as measured by one-way ANOVA comparing to siNT control. (J). Bar graphs showing results for A3B SPI-qPCR experiments. Chromatins were collected from T-47D cells transfected with indicated siRNA for 48 hours. Data represent n = 3 biological replicates and error bars for SD. * and ** denote p < 0.05 and 0.01 respectively as measured by two-tailed t-test.

To further investigate the link between TDRD3 and the binding of A3B to DNA, we conducted TDRD3 ChIP-seq in T-47D cells as well as A3B SPI-seq in cells depleted of TDRD3 by RNAi. Knockdown of TDRD3 attenuated approximately 78% of A3B binding sites identified by SPI-seq when compared to control cells, suggesting that A3B binding requires TDRD3 (**Figure 6E**). Furthermore, analysis of ChIP-seq data showed that these putative TDRD3-dependent A3B sites also displayed a strong TDRD3 binding capability (**Figure 6F-G**). We performed ChIP-seq experiments using an antibody (Millipore H3^R17me2a^) that recognises a spectrum of asymmetric dimethylarginine (ADMA)-modified proteins which are validated TDRD3 interactors^55^, and found these proteins co-localised with A3B binding sites identified by SPI-seq (**Figure 6F-G**). These results suggest that ADMA-modified proteins bind A3B and regulate its binding to DNA. When compared to DRIP-seq and DSBCapture results, a majority (62%) of the putative TDRD3-dependent A3B binding sites are R-loop-proximal, and notably, these sites also tend to overlap A3B-dependent DSBs (**Figure 6F**). From the perspective of TDRD3 binding sites, a large proportion (79%) of the *bona fide* binding sites identified using ChIP-seq overlapped with A3B binding sites, the signals of which were also attenuated by TDRD3 knockdown (**Figure S4B**). Also, more than half of the A3B-overlapping TDRD3 binding sites were proximal to R-loops, showing a bias towards A3B-dependent DSBs, and had a similar ChromHMM profile to A3B-dependent DSBs (**Figure S4B-C**). Detailed quantitative studies on seven putative TDRD3-dependent A3B binding sites (**Figure 6H**) using DRIP-qPCR and SPI-qPCR on T-47D cells confirmed a requirement of intrinsic TDRD3 on A3B binding, while depletion of TDRD3 has minimal impact on R-loop formation on these sites (**Figure 6I-J**). Therefore, our results suggests that TDRD3 may play an important role to guide the binding of A3B to regulatory elements and may serve as a mechanism to regulate its function in converting R-loops into DSBs during regulation of transcription.

Taken together, our results support a mechanistic model (**Figure 7**) in which TDRD3 functions as an adapter protein directing the DNA deamination activity of A3B to ssDNA strand of R-loops upon transactivation of ER, which also induces ADMA post-translational modifications of protein complexes localising to enhancers or promoters. The induction of co-transcriptional R-loops and A3B editing upon ER activation leads to ER-induced DNA DSBs, which modulate the chromatin configuration of enhancer or promoter regions and contribute to chromosomal instability in ER-driven cancers.

**Figure 7:**
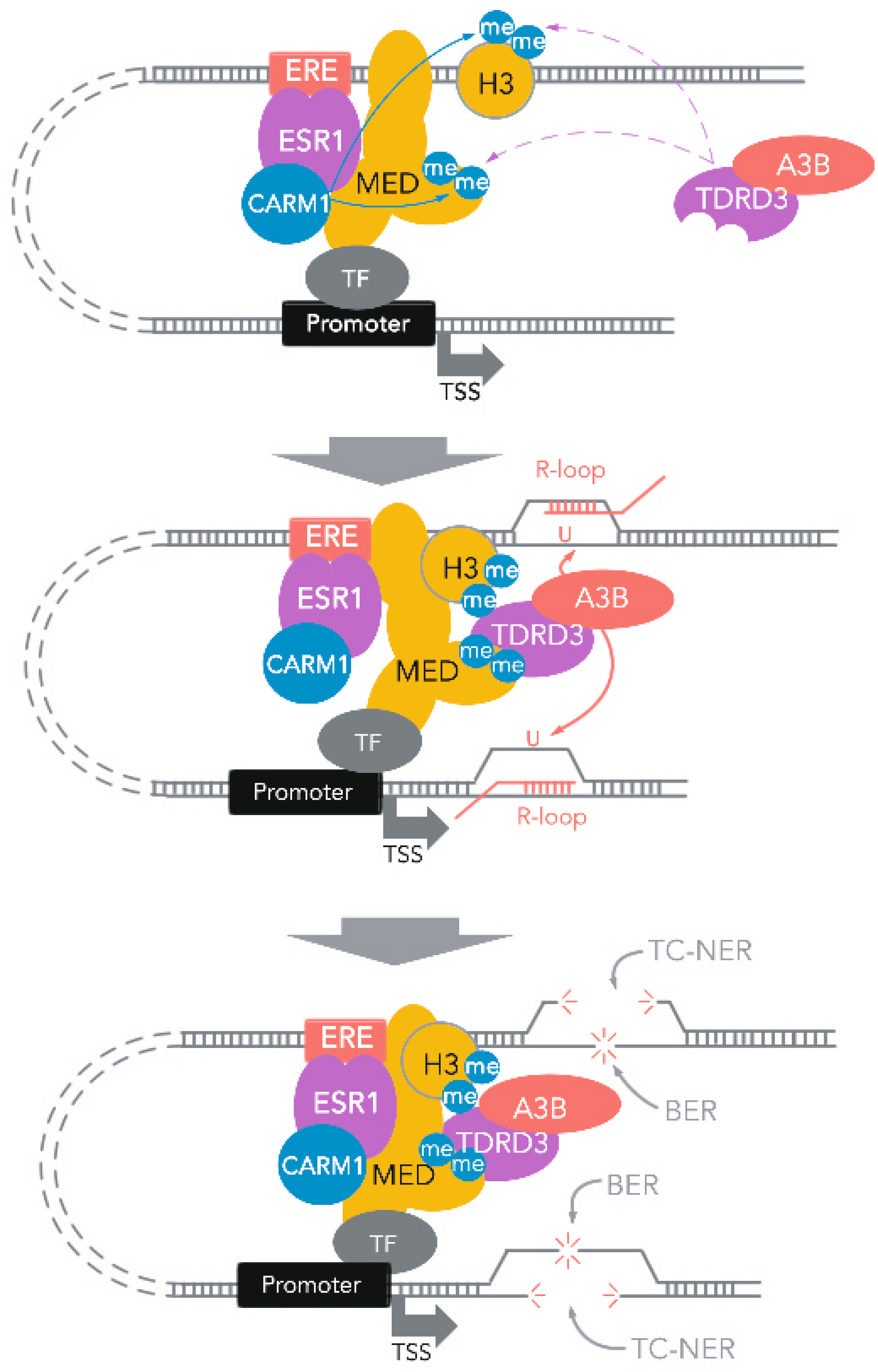
Mechanistic model detailing the generation of DSBs as a result of collaborative processing of R-loops and R-loop-editing by A3B, guided by TDRD3 adapter protein upon ER transactivation.

## Discussion

Thus far, numerous endeavours have been attempted to detect, sample and measure A3B-driven DNA deamination sites, advancing our understanding of A3B function in cancer^10, 30, 32–34^. However, direct capturing and mapping of A3B sites in cancer cells has so far proven to be challenging, mainly because of the masking effect of the highly effective BER and a TP53-dependent synthetic lethal interaction between BER-deficiency and A3B enzymatic activity^31, 36^. Here, we circumvented this problem by using a BER-deficient, A3B-overexpressing human breast cancer cell line model in a *TP53* functional loss context. By sequencing a number of clones, our approach captured A3B-driven mutations in bulk, as supported by strand-coordinated focal TC>TT hypermutations and NMF signatures with known A3B traits. The strong association between these sites with regulatory elements (TSS and enhancers), transcriptional activity and early DNA replication timing strengthens the role of A3B as a TF (in our case, ER) co-activator. We encourage additional studies beyond ER-driven models to examine whether A3B could also have a similar effect on other TFs, especially other nuclear receptors.

Using the A3B deamination sites detected by WGS, we also added R-loops into the known repository of A3B substrates in addition to lagging DNA synthesis strand^33, 35, 56^, loop regions of DNA hairpins^10, 57^, and ssDNA intermediates during recombination and repair^9, 10, 58^. The predominant pathway processing A3B-catalysed uracil lesions in the context of R-loops is the BER^24^, which has been observed in previous studies in APOBEC-Cas9-mediated R-loop editing and the AID-catalysed deamination within R-loops during immunoglobulin class switch recombination^27, 28^. However, if R-loops are resolved before repair of the uracil lesion, the re-annealing of the template strand will lead to a U:G mismatch which can also be repaired by DNA mismatch repair (MMR). Unlike the BER that faithfully repairs the U:G lesion, MMR is known to cause omikli mutations and is detectable using datasets from TCGA project and ICGC consortium^59, 60^. Nevertheless, DNA sequences flanking putative A3B-driven omikli mutations detected in these real-world datasets did not display any R-loop traits, suggesting that A3B-catalysed uracil bases in R-loops were predominantly repaired by BER.

Importantly, we also demonstrated that the repair of A3B-catalysed uracil bases in R-loops is part of a mechanism leading to the formation of R-loop-mediated DSBs, which are known to affect chromosome integrity and contribute to genomic heterogeneity in cancer cells^47, 61^. It has been shown previously that R-loop clearance by TC-NER factors, such as XPF/XPG enzymes, removes stretches of DNA/RNA hybrid and results in SSBs^48^. Further collision with the replication fork/transcriptional machinery, or actions by unidentified structure-specific endonucleases are required to generate DSBs^47, 48, 61^. Here, we have identified a unique way of R-loop-mediated DSB formation, which does not require the above-mentioned prerequisites but is achieved instead by the introduction of SSBs by BER-processing of A3B-deaminated bases at the opposite strand of R-loop-induced SSBs. This mechanism further details the role of A3B in inducing DSBs at ER binding sites^6^, and our observations unify the roles of R-loops and A3B during the formation of DSB following ER activation^6, 7^.

A fuller understanding of how SSBs effect R-loop homeostasis will require further investigation. Recently, McCann et al. demonstrated that a high-level expression of A3B in cells results in an accelerated clearance rate of steady-state- or stimulation-induced-R-loops^29^. We speculate that this process would also be accompanied by an elevated level of DNA DSBs as many of the A3B-dependent DSBs display a requirement for R-loop formation in our study. More sophisticated studies will be needed to determine whether A3B-mediated SSBs occur prior to the clearance of DNA:RNA hybrid, and if so, whether R-loops with a SSB would serve as a better substrate for TC-NER.

Using BioID-based mass-spectroscopy proteomics, we identified TDRD3 as a binding partner leading to the recruitment of A3B to enhancer/TSS R-loops. TDRD3 is an oncogene and a transcriptional coactivator^62, 63^, and can be recruited to active chromatin by the binding of its Tudor domain to ADMA-modified proteins^64^. TDRD3 exerts its function by acting as an adapter, linking these ADMA-modified proteins to effector proteins, such as TOP3B and DHX9 that resolve R-loops^54, 65^. Despite both being R-loop processors, A3B may have distinct functions compared with TOP3B, that may lead to different transcriptional outcomes. This hypothesis is supported by our observation that A3B and TOP3B compete for the same binding site on TDRD3. Previous studies have shown that loss of either TOP3B or A3B could result in R-loop accumulation and elevated DNA damage^29^. Therefore, it will be interesting to compare the DSB-inducing capabilities of these two enzymes in future studies and identify mechanisms leading to the selection of TDRD3 partners.

Our results also demonstrate that R-loop selection by A3B may be achieved by binding of TDRD3 to ADMA-modified proteins including histones (H3^R17me2a^/H4^R3me2a^), Pol II (Pol II^R1810me2a^) and the mediator complex subunit 12 (MED12^R1899me2a^)^66^. These ADMA-modified proteins mark promoter, enhancer, or looping regions, which harbour bi-directional transcriptional activity and could form R-loops with long intergenic noncoding RNA (lincRNA) or enhancer RNA (eRNA) binding^67^. Using hUGI as a surrogate of functional-loss of A3B and transcriptional profiling by RNA-seq, we showed that ER-mediated transactivation programmes rely on A3B-mediated DNA editing in these regions, consolidating A3B as a regulator of ER activity independent of ER-ligand associations.

Previous reports have proposed A3B as a therapeutic target in a model where elevated A3B drives tamoxifen-resistance through the acquisition of increased A3B-dependent gene mutations. Our data also support the therapeutic targeting of A3B in tamoxifen-resistant breast cancer but with an emphasis on its role as a driver of an adaptive response at the transcriptional or epigenetic level rather than through direct gene mutation. Our findings also support the development of A3B inhibitors as new therapies to control ER activity in the context of selective ER modulator-resistant cancers and, potentially, additional nuclear hormone receptor-driven malignancies.

## Materials and methods

### Cell culturing and cell line preparation

T-47D (ATCC HTB-133), MCF-10A (ATCC CRL-10317) and MDA-MB-468 (HTB-132) cells were obtained from ATCC. Lenti-X 293T cells were obtained from Clonetech. Cells were maintained using ATCC recommended medium except Lenti-X 293T cells were cultured with DMEM medium supplemented with 10% v/v FBS (PAA) and 0.5% v/v 10000U/ml penicillin /streptomycin solution. All cells were cultured at 37 °C with 5% CO_2_.

To generate stable inducible cell lines, T-47D human breast cancer cells at exponential grow phase were transfected with lentivirus at a M.O.I of 100 with the assist of 10 μg/ml polybrene. After a transduction period of 48 hours, selection was carried out using medium supplemented with 4 μg/mL puromycin (Gibco). Transduction efficiency was monitored by either turbo-RFP expression, A3B immuno-fluorescence microscopy or immunoblotting following induction of sample cells with doxycycline. After verification, lentivirus cassettes were maintained by culturing cells with 4 ug/mL puromycin.

### Drug treatments

For the induction of lentiviral protein expression, cells were pre-exposed to full culture medium supplemented with 1 μg/mL doxycycline (Sigma) for 24 hours unless denoted otherwise. Follow-up drug treatments, if included, also contained the same concentration of doxycycline throughout the entire treatment period.

For ER induction by E2 (Selleck Chem), adherent cells were briefly washed by warm PBS twice before incubated with phenol-red-free RPMI-1640 medium (Gibco) supplemented with 10% charcoal-stripped FBS (Thermo Fisher) for 48 hours at 37 °C. E2 was added to the culture medium at a final concentration of 100 nM, and further incubated for two hours (unless denoted otherwise) before the cells were harvested for follow-up studies.

### Colony separation and WGS

0.25 million exponentially growing T-47D cells harbouring lentiviral 3×Flag-A3B-P2A-hUGI-HA or 3×Flag-A3B**-P2A-hUGI-HA were seeded onto a 100 mm plate for 24 hours, followed by induction with 1 μg/ml doxycycline for 120 hours. Cells were harvested using trypsin digestion, serial diluted and plated onto a 96-well plate (Corning). Colonies derived from single cells were picked and the cloned cells were expanded with culture medium supplemented with 4 μg/ml puromycin. Genomic DNA was extracted from each colony using a Qiagen Genomic-tips 500/G kit. Sequencing libraries for whole genome sequencing with DNA nanoball (DNB) technology were constructed (Beijing Genomics Institute (BGI) Inc., Hong Kong). Sequencing was carried out using BGISEQ-500 sequencer with a mean sequencing coverage of greater than 30× for each of the samples using 2 × 150 bp configuration.

Further method details are provided in the supplementary text.

## Data availability

WGS, ChIP-seq, SPI-seq, ssDRIP-seq, DSBCapture-seq and RNA-seq datasets reported in this study are available as dataset GSE193234 in the Gene Expression Omnibus (GEO) database. Processed results for RNA-seq are available in Supplemental Table 1, and processed BioID mass spectrometry are available in Supplemental Table 2. Previously reported datasets used in this study are listed in the supplemental text.

## Acknowledgements

This work is financially supported by Cancer Research UK (C309/A11566 and C2739/A22897) and ICR (London, United Kingdom). P.W. acknowledges additional grant support from the Wellcome Trust (212969/Z/18/Z), Cancer Research UK (C35696/A23187) and The Mark Foundation/The Chordoma Foundation. C.Z. was sponsored by Shanghai Pujiang Programme (22PJD104). We thank Prof. Ping Yuan from Sun Yat-sen University, Dr. Mike Walton and Dr. Alexandra Vasile from ICR for helpful discussions.

## Author Contributions

C.Z., O.W.R., P.A.C, and P.W. conceived and designed this study. C.Z., Y.L., M.W., and M.T. conducted the experiments and analysis unless otherwise noted. B.C.,Z.B., K.M. and B.A-L conducted analysis on sequencing data. provided guidance on data analysis using sequencing data. C.Z., A.H., M.P.L, O.W.R., P.A.C, and P.W. prepared the manuscript.

