## Supplemental Materials for "R-loop editing by DNA cytosine deaminase APOBEC3B determines the activity of estrogen receptor enhancers"

**SI contents:**

1. Supplemental figure S1-S3, and legends
2. Supplemental methods
3. Supplemental reference
4. Supplemental table S1-S2

### Supplemental Figures

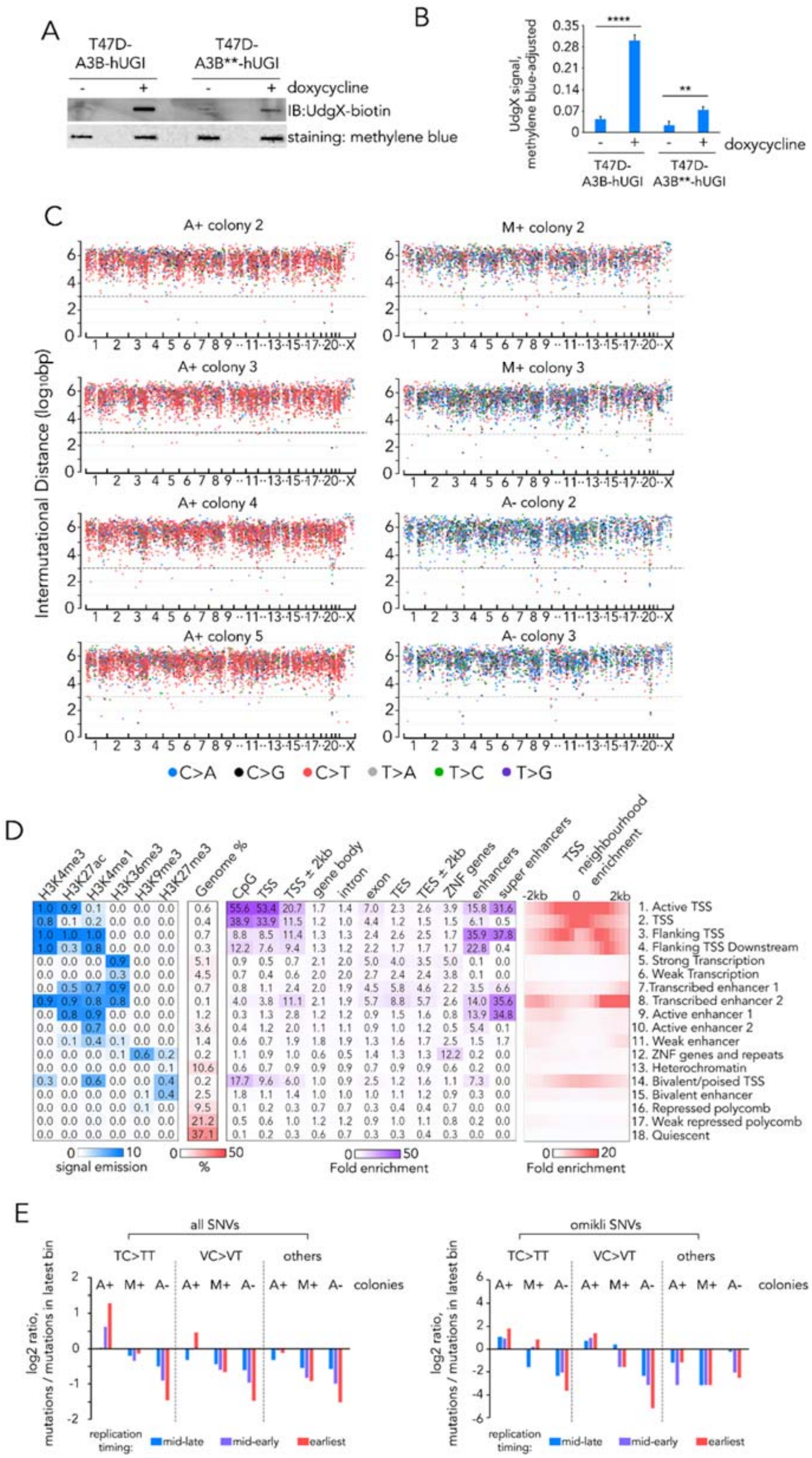

**Figure S1:** Capture and characterisation of A3B editing sites in BER-deficient cell models (related to Figure 1).

(A). Representative slot blots of genomic DNA extracted from indicated cells. Uracil signal was detected using biotinylated UdgX protein and total DNA was detected using methylene blue staining.

(B). Level of uracil incorporation for indicated cells measured by adjusted UdgX slot blot signals. Data represent average values from 3 biological replicates and error bars for standard deviation. ****: $p \leq 10^{-4}$; **: $p \leq 0.01$, two-tailed Student's T-test.

(C). Waterfall plot of intermutational distance (IMD) of each mutation identified in the indicated colonies. Dotted line denotes IMD ≤ $10^3$ bp.

(D). The 18-state ChromHMM model used in this study was derived from six epigenetic marks. For validation, enrichment scores for ChromHMM-curated known genomic regions as well as enrichment scores for TSS neighbourhood regions are shown. In addition, enhancer segments from Enhancer Atlas[1] and super enhancer segments for T-47D cells[2] were included for the enrichment analysis.

(E). Rates of indicated type of mutations in replication timing quartiles, relative to the latest replicating quartile, for SNVs from indicated sample groups.

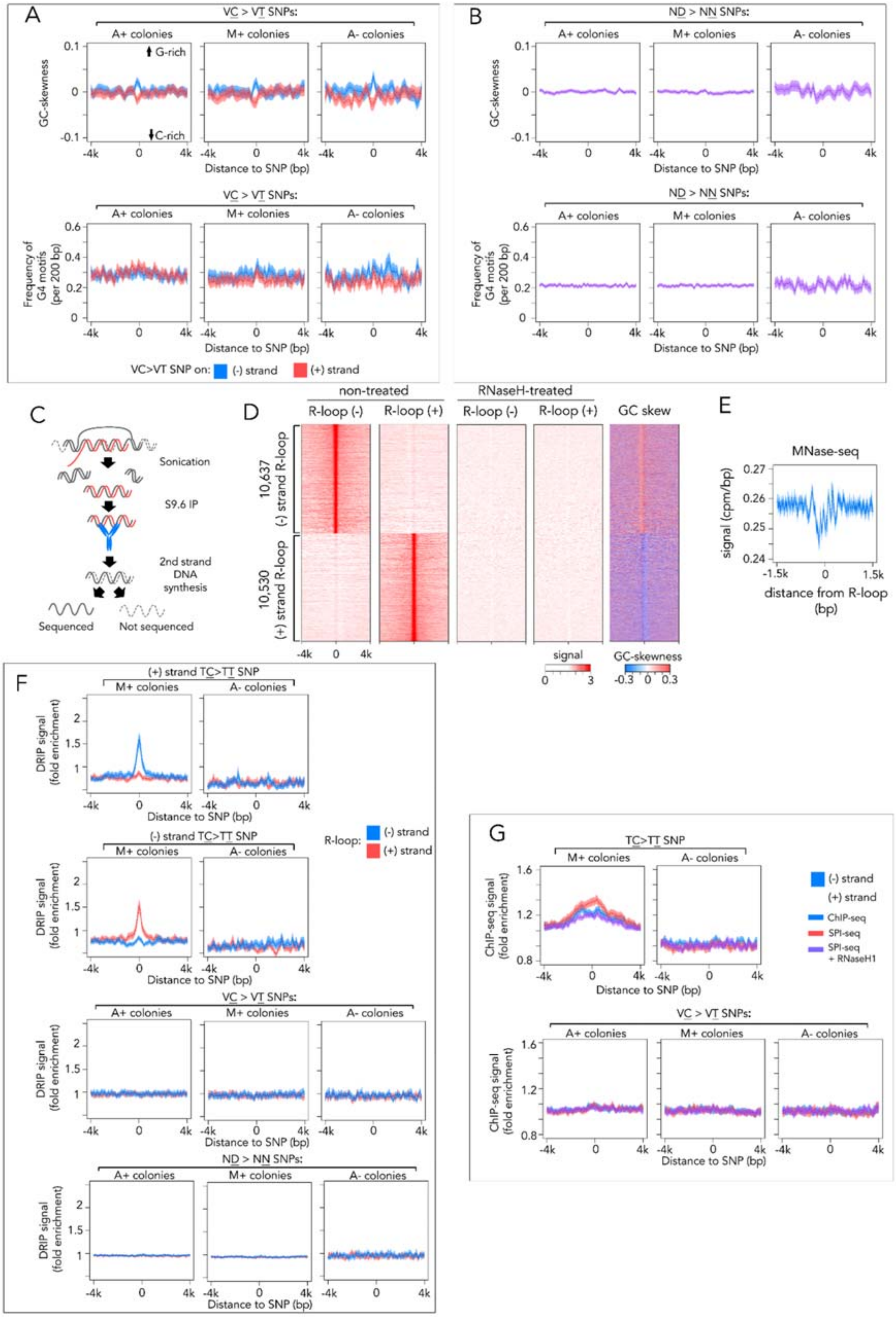
A
VC > VT SNPs:
A+ colonies
M+ colonies
A- colonies
GC-skewness
G-rich
C-rich
Distance to SNP (bp)
Frequency of G4 motifs (per 200 bp)
VC>VT SNP on: (-) strand (+) strand
B
ND > NN SNPs:
C
Sonication
S9.6 IP
2nd strand DNA synthesis
Sequenced
Not sequenced
D
non-treated
RNaseH-treated
R-loop (-)
R-loop (+)
GC skew
10,637 (-) strand R-loop
10,530 (+) strand R-loop
signal
GC-skewness
E
MNase-seq
signal (cpm/bp)
distance from R-loop (bp)
F
(+) strand TC>TT SNP
(-) strand TC>TT SNP
DRIP signal (fold enrichment)
R-loop: (-) strand (+) strand
G
TC>TT SNP
ChIP-seq signal (fold enrichment)
ChIP-seq
SPI-seq
SPI-seq + RNaseH1

**Figure S2:** Mapping R-loops in T-47D genome using strand-specific DRIP-seq (ssDRIP-seq, related to Figure 2).

(A). Profiles of GC skewness and frequency of G-quadruplex (G4) motifs in regions flanking the VC>VT SNP identified in indicated colonies.

(B). Profiles of GC skewness and frequency of G-quadruplex (G4) motifs in regions flanking the ND>NN SNP identified in indicated colonies.

(C). Schematic of the ssDRIP-seq experiment carried out in this study.

(D). Heat maps of signal from ssDRIP-seq and GC skew score in regions flanking high-confidence R-loops identified by MACS2 programme.

(E). Profile of MNase-seq signals in region flanking centre of R-loops. Line denotes average value and shaded areas 95% CI.

(F). Signal profiles showing ssDRIP-seq signals in regions flanking the TC>TT SNP or VC>VT SNP identified in indicated colonies on indicated strand.

(G). Signal profiles for Flag-A3B ChIP-seq or SPI-seq signal in regions flanking TC>TT SNP or VC>VT SNP identified in indicated colonies

For A-B, and E-G, lines denote average value and shaded areas 95% CI. Rolling windows with size of 200 bp was used to calculate GC skew score and G4 motif frequency.

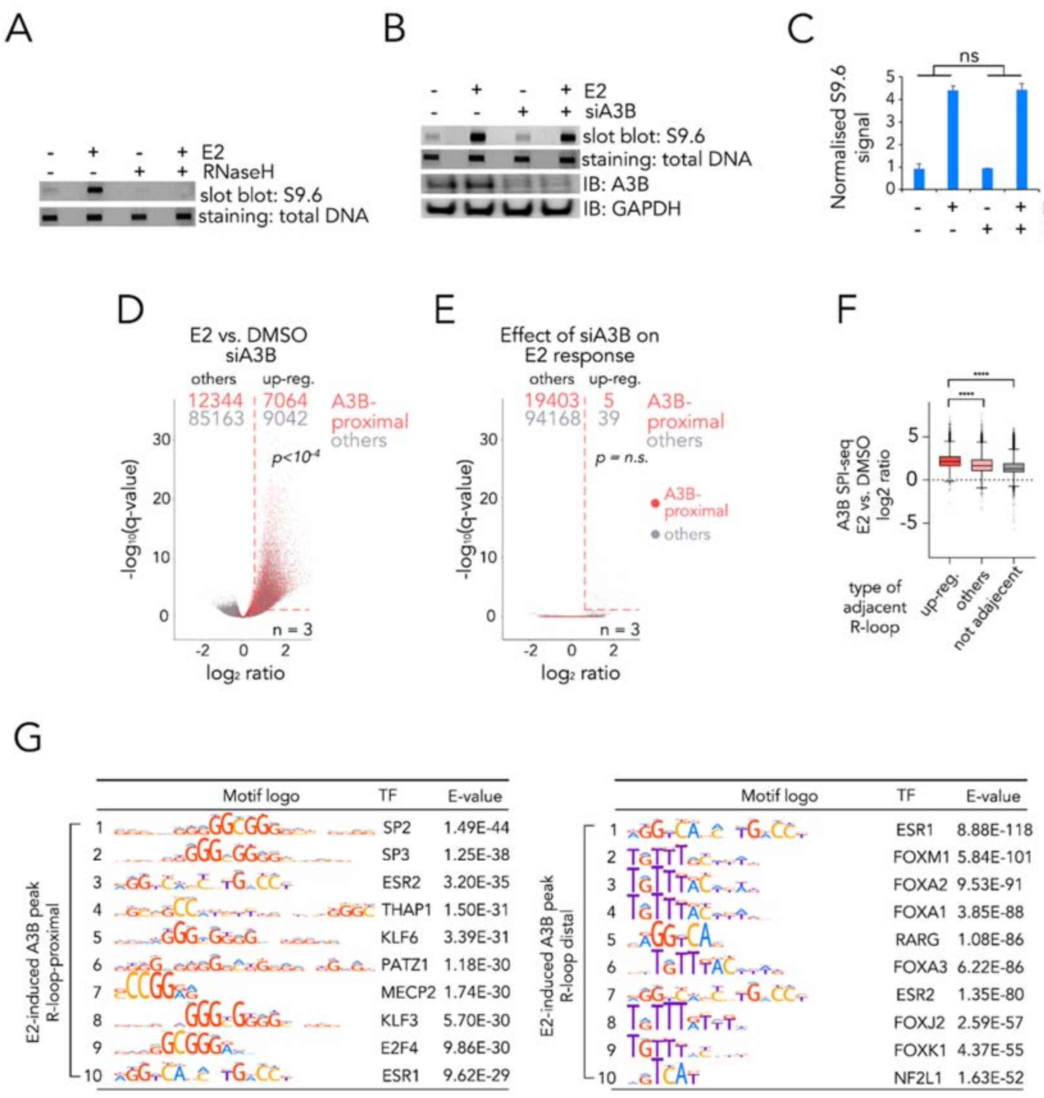


**Figure S3:** E2-induced R-loop formation was not dependent on A3B but might alter A3B binding pattern (related to Figure 3).

(A). Representative slot blots of gDNA samples from T-47D cells with or without two-hour stimulation by 100 nM E2. To confirm specificity of S9.6 antibody, gDNA samples treated overnight with RNaseH were included as control.

(B). Representative slot blots and immunoblots of samples from T-47D cells treated with or without two-hour stimulation by 100 nM E2, and with or without depletion of A3B by siRNA.

(C). Bar graph depicting data analysis of B. Data represent average value of methylene blue-adjusted S9.6 signal from two biological repeats (error bars indicate standard deviation). NS denotes non-significant from two-way ANOVA assessing size effect of A3B depletion on E2 response.

(D). Volcano plot summarising the effect of two-hour stimulation by E2 on R-loop formation in A3B siRNA-treated cells, determined by ssDRIP-seq using DESeq2 statistics. p value represent statistical significance by $\chi^2$ test of non-random association between R-loop or A3B proximity and response to estradiol. Red dotted lines show region meeting hit criteria.

(E). Volcano plot summarising the effect of A3B depletion on E2 response of R-loop level, determined by ssDRIP-seq using DESeq2 statistics. For D and E, A3B-proximal R-loops are coloured in red. Criteria for up-regulated R-loops are defined as FDR $\leq 0.05$ with fold change $\geq 1.5$ and is labelled with red dotted line. p value represent statistical significance by $\chi^2$ test of non-random association between R-loop or A3B proximity and response to estradiol.

(F). Tukey boxplots showing log2 ratio of SPI-seq signals for Flag-A3B binding sites in response to estradiol. Data derived from two biological replicates for each condition using edgeR. ****: $p \leq 10^{-4}$, one-way ANOVA.

(G). Top ranking transcription factor motifs enriched in R-loop-proximal (left) or R-loop-distal (right) Flag-A3B sites that were induced by E2. Enrichment scores (E-values) were derived by AME from MEME suit using shuffled input sequences as background control.

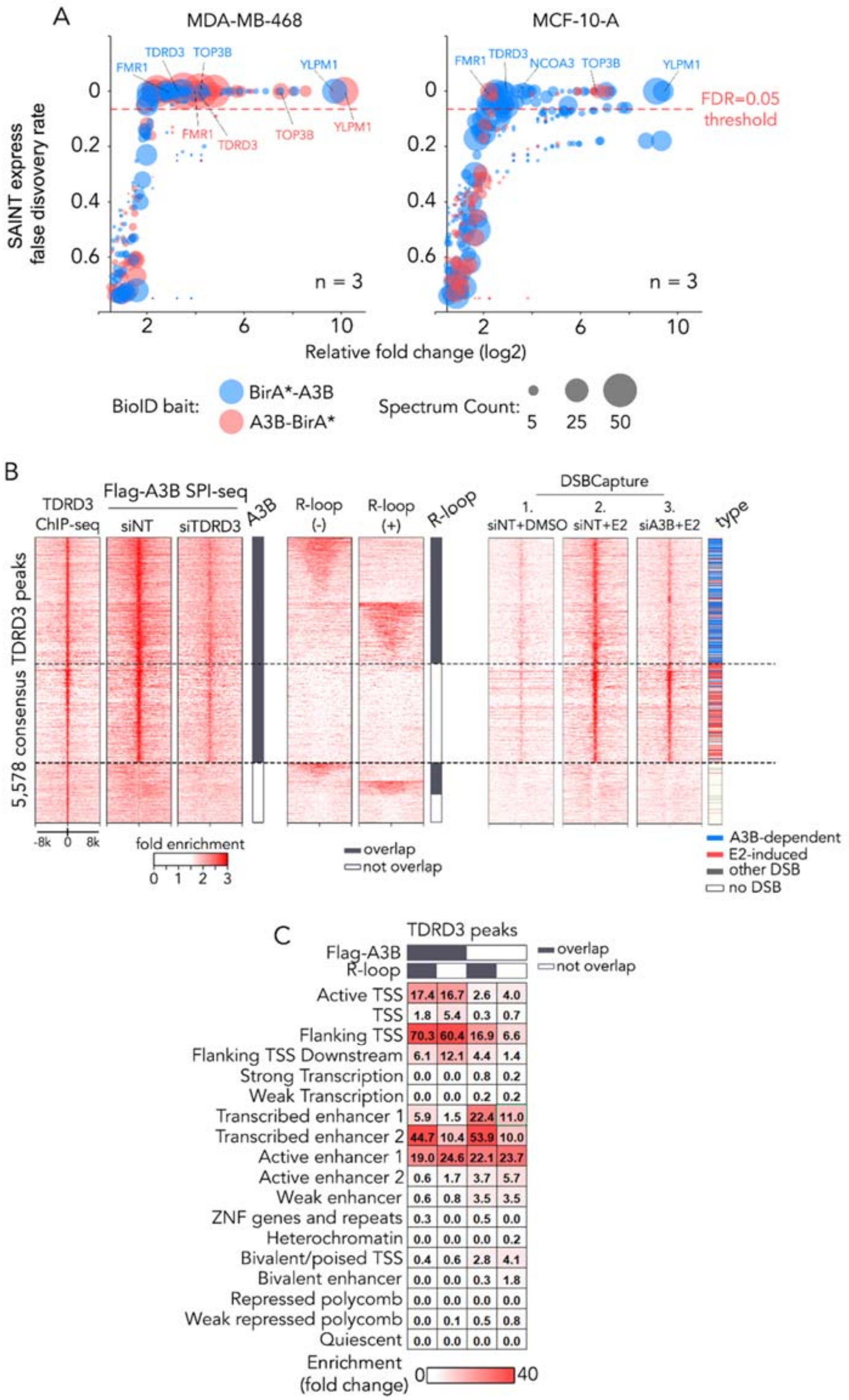


**Figure S4**:TDRD3 interacts with A3B and guides its binding.

(A). Bubble plot illustrating the result of BioID experiments for MDA-MB-468 and MCF-10-A cells. Cells bearing lentiviral inducible BioID baits were induced with doxycycline for

24 hours before subject to streptavidin affinity purification and mass-spec. SAINT express FDR score were derived using BirA*-RFP pull-down results as control and calculated by CRAPOME. Data derived from n = 3 independent experiments and recurring hits were labelled.

(B). Heat maps illustrating signals from indicated sequencing experiments in ±8 kb regions flanking TDRD3 binding sites identified by TDRD3 ChIP-seq experiments. Also shown are bars indicating whether TDRD3 sites are overlapping A3B binding sites or R-loops, as well as the types of overlapping DSBCapture peaks.

(C). Heat map showing enrichment scores for ChromHMM chromatin states at indicated TDRD3 binding sites.

#### Supplemental Methods

##### Plasmids cloning and lentivirus packaging

DNA encoding 3×Flag-A3B-P2A-hUGI-HA were synthesised by GeneArt service (Thermo Fisher Scientific Inc.). In order to distinguish the mRNA of ectopic-expressed proteins from the intrinsic ones and boost the expression, codons for both A3B and hUGI were optimised using GeneArt algorithm.  AgeI restriction site and Kozak sequences were introduced at the 5'-end and BspD1 restriction site was introduced at the 3'-end of the construct using PCR method. Empty pTRIPZ plasmid was obtained from Horizon Discovery. The DNA fragment was introduced to pTRIPZ plasmid using AgeI and BspD1 restriction digestion and ligation methods with enzymes provided by New England Biolabs. For A3B** construct, two rounds of site-directed mutagenesis at E68Q and E225Q sites on A3B were performed on 3×Flag-A3B-P2A-hUGI-HA-containding pTRIPZ plasmid using the Quick-Change kit provided by Agilent. The resultant plasmids were sequenced and maintained in Mach1 T1 E. coli cells.

DNA encoding codon-optimised A3B and turbo-RFP were synthesised by GeneArt service (Thermo Fisher Scientific Inc.) and cloned into gateway entry plasmids. The destination plasmids from a previous study[3], namely pDEST_pcDNA5_BirA-FLAG_Nterm and pDEST_pcDNA5_BirA-FLAG_Cterm was kindly supplied by Dr. Anne-Claude Gingras from

Lunenfeld-Tanenbaum Research Institute, Toronto. After cloning A3B and turbo-RFP into the two destination plasmids (resulting plasmids encoding BirA*-A3B, A3B-BirA* and BirA*-RFP) using gateway technology and enzymes (Invitrogen Inc.), the fragments were cloned into pTRIPZ plasmid at AgeI and BspD1 site. The resultant plasmids were sequenced and maintained in Mach1 T1 *E.coli* cells.

For lentivirus packaging, helper plasmids, namely psPAX2 and pMD2.G were co transfected with pTRIPZ encoding protein of interest into Lenti-X 293T cells (Clonetech) using calcium-phosphate transfection kit (Promega). After changing the culture medium after 24 hours, virus was harvested at 48 and 72 hours, pooled, precipitated using Peg-it reagent (System Biosciences) and resuspended in serum-free RPMI medium (Gibco). Multiplicity of infection (M.O.I) of viruses were measured using manufacture's guidance (Horizon Discovery).

**Immunoblotting**

Cells were lysed in RIPA buffer (Millipore) containing complete protease inhibitor cocktail (Roche) and phosphatase inhibitors solution 2 (Sigma). Protein concentrations were measured by BCA protein assay (Pierce). Samples were separated on SDS polyacrylamide gels and transferred to PVDF membranes using the iBlot2 system (Invitrogen). Membranes were washed with TBST (25 mM Tris, 140 mM NaCl, 0.1% Tween-20, pH 7.5) and probed with various primary antibodies overnight at 4ºC, followed by binding of IRDye 680- or IRDye 800CW-conjugated secondary antibodies (LI-COR) for 1 hour at room temperature. The imaging and quantification of protein bands was carried out using an LI-COR CLX infrared imaging system.

**UdgX preparation and slot blotting**

DNA encoding M. smegmatis UdgX protein with N-terminal 6×His-tag and C-terminal AviTag was synthesised by GeneArt (Thermo Fisher) and cloned into pET24a plasmid. Biotinylated UdgX protein was expressed in AVB101 E. coli strain, and sequentially purified using Ni-NTA and Superdex75 columns on an ÄKTA Avant protein purifier.

Uracil-incorporated positive control DNA was obtained by extraction of pUC19 plasmid which was transformed into K12 CJ236 E. coli strain (NEB). Genomic DNA from T-47D cells was extracted and purified using Qiagen Genomic-tips 500/G kit following manufacturer's instructions. Sample DNA (100 ng for positive control and 4 µg for genomic DNA) was loaded onto a PVDF membrane with Hoefer PR600 slot blot manifold linked to a vacuum pump. The membrane was blocked by Pierce protein-free blocking buffer (37572), followed by incubating with 1 µg/ml biotinylated UdgX at room temperature for 1 hour. After probing with streptavidin-HRB conjugate (Pierce), the membranes were washed with TBST and detected with LI-COR CLX infrared imaging system using ECL-plus substrate (Pierce).

**SNV calling and analysis**

Raw sequences from BGISEQ-500 sequencer were aligned to the human genome (hg38) Burrows–Wheeler Aligner (BWA)[4]. The resultant bam files were then recalibrated using GATK 4.0.5 mutation calling kit following the 'best practice' advised by the developers[5]. Briefly, bam reads were reordered, marked with group and duplication status with Picard tools, followed by base quality score recalibration by GATK BaseRecalibrator programme. Joint variant calling was performed using GATK Mutect2 programme, with doxycycline-induced colonies as 'Tumour' sample and non-induced colonies as 'Normal' sample. In order to improve mutation calling accuracy, resources common germline variant sites from Genome Aggregation Database (gnomAD) and a 'panel-of-normal' dataset consisting of several wild-type T-47D mutation calling results were included in the Mutect2 analysis[6]. After calculating the potential sampling contaminations using GATK CalculateContamination programme, the Mutect2-derived SNVs were then selected against 14 filters implemented by the GATK FilterMutectCalls. In order to exclude SNVs that are located on either ends of homopolymer DNA sequences and within simple repeating elements, both of which are likely to generate false-positive results, the SNVs passing through GATK filters were further filtered using SNPiR software[7] with perl scripts filter_homopolymer_nucleotides.pl and pblat_candidates_ln.pl scripts only. The latter script was enabled by pBLAT algorithm[8] and BEDTools programme[9].

The resultant SNVs, encoded in vcf file format, were subjected to further analyses using programmes coded by R language (R Core Team). Calculation and visualisation of intermutational distance by waterfall plot was done with R package ‘qqman’, with minor modifications on source codes[10]. Kataegis and omikli mutation clusters were detected using the hyperclust programme, with the methods described previously[11]. The effect of variants was analysed using Variant Effect Predictor (VEP)[12], with the assistance of genome assembly lift over tool provided by UCSC Genome Browser[13]. Random sampling and enrichment analysis by Fisher exact test for variant effects were conducted using bedtools. For replication timing analysis for SNVs, a method described previously[11] was employed using T-47D Repli-seq dataset from ENCODE (ENCFF440QFG).

**Mutational signature analysis**

All SNVs detected in the M+，A+ and A- groups were used for identifying mutational signature contributions. Mutation signatures and spectrum analysis were analyzed by Bioconductor package MutationalPatterns with 30 COSMIC signatures following the standard workflow[14]. *De novo* mutational signature extraction was performed using the NMF algorithm[15]. Cosine similarities between *de novo* extracted signatures and the COSMIC cancer mutational signatures were also calculated with MutationalPatterns. Absolute and relative contributions of each de novo obtained signature were also determined.

**Chromatin state analysis**

Chromatin state characterisation was conducted using ChromHMM software[16]. To provide modelling input, raw reads from ChIP-seq results using six chromatin marks were downloaded from GEO database and aligned to GRCh37 (hg19) genome assembly using BWA. The resultant bam files were binarised and used as training set to build the Hidden Markov Model that segments and annotates T-47D genome. To validate the chromatin model generated, degree of overlap between known genomic features and model was carried out using script in ChromHMM. The genomic features were pre-defined by ChromHMM authors, except that the super enhancer regions in T-47D cells were derived as previously reported[17]. For computing the overlap between ChromHMM and genomic regions of interest, the full-length peak regions were used except for SNVs where ±200 bp flanking regions were used.

**Strand-specific RNA/DNA immunoprecipitation (ssDRIP) and sequencing**

RNA/DNA immunoprecipitation was performed on T-47D cells using an optimised method previously reported by Halász et al[18]. The entire procedure followed the best-performing experiment number 5 in the report, the detailed protocol of which was provided in the supplementary information section, except that Covaris E220 was used for fragmenting genomic DNA with the following settings: duty factor 10%; peak incident power 75 watts; cycles per burst 1000; time 50 seconds; temperature 4 ºC. S9.6 antibody was purchased from Kerafast (ENH002), and mouse IgG (Thermo Fisher 31903) was used as control (if required).

ssDRIP-seq was carried out with immunoprecipitated DNA:RNA hybrids. To enable strand-specificity, the product was subjected to the second-strand DNA synthesis with uracil incorporation. More specifically, DNA:RNA hybrid was reconstituted in 1× NEB first-strand synthesis buffer (NEB E7525) without the addition of primer and enzyme to a final volume of 20 µl. The product was immediately used as the starting material of NEB's directional second-strand synthesis method following manufacturer's instructions (NEB E7550). The resultant DNA was purified using SPRI beads (Thermo Fisher) at a volume ratio of 2.0 and eluted using TE buffer. After quality inspections by BioAnalyzer 2100 (Agilent), the product was sent to BGI, Hong Kong, where strand-specific libraries were constructed using DNB technology. The resultant libraries were sequenced with BGISEQ-500 instrument.

**Data processing of ssDRIP-seq**

Pair-end raw reads were aligned to GRCh37 genome assembly using BWA, followed by file format exchange by SAMtools. The resultant alignment reads were separated by first-in-pair strand into two files corresponding to (+) or (-) strand R-loop (or input) reads using SAMtools flag identifiers. Peak calling was carried out using MACS2 package using pair-end fragment setting with input and ssDRIP samples. Signal tracks was generated using MACS2, where score represents fold enrichment over input control.

For quantification of ssDRIP experiments, consensus R-loops peaks were generated first. The resultant peaks for 12 replicates were served as input for MSPC, and recurrent peaks were filtered using parameters '–w 1E-4 –s 1E-8 –c 4'. Peaks on the same strand and within a distance of 1000 bp were merged using BEDTools and converted to GTF format using UCSC tools (bedtogenepred and genepredtogtf). Read counting was conducted using Subread, and raw counts were analysed by DEseq2 using a generalized linear model with an interaction term.

**GC skewness and G4 motif analysis**

GC-skewness was calculated as previously described. Genomic GC-skew data, which is encoded as a bigwig file format with a window length of 200 bp, was created by R-loop DB[19] and downloaded from UCSC genome browser[13].

DNA G4 motif was predicted using G4Hunter[20], which is encoded in a R package. Scan was performed using a G4 score threshold of 1.2 on GRCh37 genome assembly, followed by composing the resultant file in bigwig format with a window length of 100 bp for follow-up studies using R scripts provided by authors of G4Hunter.

**Sequencing data visualisation**

Bigwig files containing enrichment scores, GC-skewness score and G4 motif frequency across GRCh37 genome assembly was used to plot the heatmaps and profile plots. The R package 'seqplots' was used to perform data visualisation with the assistance of a graphical user interface[21]. For heatmaps, average data using a window length of 50 was used, whereas for profile plots, average data with standard deviation using a window length of 100 was used. Signal profiles across specific genomic regions were visualised by The Integrative Genomics Viewer (IGV)[22].

**S9.6 co-immunoprecipitation**

S9.6 co-immunoprecipitation was conducted using the method as previously reported[23], except that S9.6 antibody was purchased from Kerafast (ENH002) and IgG control from Thermo Fisher (31903). Immunoblotting were carried out using S9.6-co-immunoprecipitated proteins, with HRP-conjugated secondary antibodies and SuperSignal West Femto substrate (Thermo Fisher 34094) and visualised by LI-COR CLX infrared imaging system.

**RNA/DNA hybrid slot blots**

Genomic DNA from T-47D cells was extracted and purified using Qiagen Genomic-tips 500/G kit following manufacturer's instructions. 8 µl DNA was treated with or without RNase H (NEB M0297) at a final reaction volume of 400 µl overnight at 37 ºC and loaded onto a pre-wetted Hybond N+ nylon membrane (Cytiva, RPN303B) with Hoefer PR600 slot blot manifold linked to a vacuum pump. The membrane was immediately transferred to a Stratalinker UV crosslinker (model 1800 Stratagene) and crosslinked using a total energy of 120,000 microjoules. The membrane was then wetted with TBST and blocked with StartingBlock T20 blocking buffer (37543) at 4 ºC overnight, followed by probing with S9.6 antibody and anti-mouse-HRP conjugate (Cell Signalling 7076). The membrane was visualised using LI-COR CLX infrared imaging system with SuperSignal West Femto HRP substrate (Thermo Fisher 34094).

**Protein transient over-expression and siRNA transfection**

ppyCAG_RNaseH1_WT plasmid was used for transient over-expression of RNase H1 protein and was a gift from Xiang-Dong Fu (Addgene plasmid # 111906). This plasmid encodes V5-tagged human RNase H1 without the first 27 amino acids and replaced with a nuclear localisation signal. For protein transient over-expression, T-47D cells were plated in a 6-well plate at a density of $5\times10^4$ cells per well and incubated for 24 h. After brief washing with warm PBS and replacing the medium with RPMI-1640 medium supplemented with 10% charcoal-stripped FBS, 1.5 µg of the vector was transfected into the cells by using Lipofectamine LTX

PLUS (Invitrogen A12621) according to the manufacturer's instructions. Cells were then treated with DMSO or E2 and proceeded with subsequent experiments.

For RNAi experiments with siRNA transfection, the same cell culture preparation and follow-up experimental procedures were used as plasmid transfection, except that Lipofectamine RNAiMAX transfection reagent was used according to manufacturer's instructions, and plasmid DNA was replaced with 20 nM siRNA. For DSBCapture experiments, cell culture and transfection reagent were scaled up to 100 mm plates according to manufacturer's recommendations.

**Chromatin immunoprecipitation (ChIP)**

ChIP protocol used in this study was adapted from the protocol used by Myer's Lab at HudsonAlpha (protocol version v011014) which contributed to the ENCODE project[24] with the following modifications. Firstly, fixed T-47D cells were lysed in ChromaTrap lysis buffer that designed for sonication (Porvair Science, 100001), and chromatin fragmentation was carried out on a Covaris E220 instrument with optimised settings for ChIP purpose (duty factor 2%; peak incident power 105 watts; cycles per burst 200; time 20 mins; temperature 4 ºC). The resultant chromatin was digested with proteinase K (Qiagen) and DNA concentration was measured with a Qubit fluorometer (Thermo Fisher). Secondly, Pierce ChIP-grade protein A/G beads (Pierce 26162) were used to perform immuno-precipitation experiments, and binding of chromatins to beads were carried out at 4 ºC overnight. Thirdly, for different antibodies, optimised compositions of wash buffers are used and are as follows: Flag-M2 (100 mM Tris pH 7.5; 500 mM LiCl; 1% NP-40; 1% sodium deoxycholate); $H3^{R17me2a}$ and CARM1 (100 mM Tris pH 7.5; 500 mM LiCl; 1% NP-40; 0.1% sodium deoxycholate); TDRD3 (100 mM Tris pH 7.5; 500 mM LiCl; 0.5% NP-40; 0.5% sodium deoxycholate). Finally, the resultant immuno-precipitated DNA was purified by SPRI beads (Thermo Fisher) using a ratio of 2.0 and analysed using BioAnalyzer 2100 (Agilent).

**Single-strand DNA–associated protein immunoprecipitation (SPI)**

SPI was carried out following developers' literature with modifications[25]. Namely, the same method was used to capture chromatin-bound DNA as described in the previous session, except that the resultant DNA was denatured at 95 ºC and immediately quenched on ice.

**Sequencing and Data processing for ChIP/SPI-seq**

ChIP and SPI samples were sent to Beijing Genomics Institute (BGI Inc.) for library construction and sequencing. For ChIP samples, a standard pipeline for the construction of DNB library was used, whereas for SPI samples, the strand-specific DNB library construction protocol was requested in order to include ssDNA species in the library. Sequencing was carried out on a BGI-500 instrument with single-end 50 bp read length for both input and immuno-precipitated DNA.

For data analysis, raw reads were aligned to GRCh37 genome assembly using BWA, followed by file format exchange by SAMtools[26]. Peak calling was carried out using MACS2[27] package using paired input/IP sequences with modelled narrow peak settings and an extended fragment length of 200 bp. Enrichment score was computed by MACS2 using the resultant files from peak calling and resented as fold change of IP over control. For n=2 replicated ChIP/SPI-seq data (TDRD3 ChIP-seq), reproducible peaks were picked using MSPC[28].

Peaks were filtering against a 'blacklist' region defined by the ENCODE project[24]. For quantitative analysis of the effect of E2 over DMSO on SPI-seq data, EdgeR[29] statistics were performed with the R package Diffbind[30], using narrow peak produced by MACS2 and aligned sequence reads. To filter for peak of interest, top 25% peaks by base mean score from normalised sequencing read counts were included for the down-stream analyses. Significant peaks were defined as FDR $\leq 0.05$ with fold change $\geq 1.5$ by Diffbind. Intersection between peak regions and Fisher's exact test with randomised genomic regions were performed by BEDTools, and data was visualised by the R package SeqPlots. Transcription factor binding motif analysis was carried out using the AME[31] programme from the MEME[32] programme suite. Distance between peak features were analysed by BEDTools[9], where proximity was defined as ± 1.5 kb of distance.

**DSBCapture-seq**

DSBCapture was conducted following a previous report[33] except that Covaris E220 was used for fragmenting adapter-ligated DNA with the following settings: duty factor 10%; peak incident power 75 watts; cycles per burst 1000; time 50 seconds; temperature 4 ºC.

**Data processing for DSBCapture-seq**

DNA libraries generated by DSBCapture protocol were sequenced using Illumina HiSeq2500 by BGI. Raw reads were aligned to GRCh37 genome assembly using BWA and processed by SAMtools. MACS2 was used to perform peak calling, with modelled narrow peak settings and an extended fragment length of 200 bp. Signal tracks were generated by MACS2, where score represents fold enrichment over input control. For quantification, EdgeR analysis was performed using Diffbind using the same procedure as ChIP-seq. Heatmap was created using R package ComplexHeatMap[34].

**RNA-seq and quantification**

Total RNA from T-47D cells were extracted using a MagNA pure 96 instrument (Roche), and RNA integrity number (RIN) was determined by BioAnalyzer 2100 (Agilent). RNA libraries were constructed and sequenced by BGI using poly-dT enrichment in conjunction with DNB technology on BGISEQ-500 instrument. Raw reads were aligned to GRCh38 genome assembly and GENCODE[35] GRCh38.p13 annotation using STAR programme[36], followed by processing of the resultant file using SAMtools. Read counting on genomic features was carried out using Rsubread[37], and the R package DESeq2[38] was used to perform statistic-based quantification. In DESeq2 analysis, in addition to terms to denote doxycycline and E2 effects in the design formula, an additional interaction term was added to denote the synergism effect between the two drugs following the user's manual written by the DESeq2 authors (http://www.bioconductor.org/packages/devel/bioc/vignettes/DESeq2/inst/doc/DESeq2.html). Expressed genes in T-47D cells were defined as base-mean of normalised reads greater than 50% of all genes. Differentially expressed genes were picked using an FDR cut-off of 0.05 and fold change cut-off of 1.5. Heatmap was created using ComplexHeatMap.

**Gene set analysis**

Gene set analysis was carried out using MSigDB[39, 40] with gene set collection H using differentially expressed gene identified by RNA-seq. Cis-regulatory region-associated genes were predicted using rGREAT[41] using differential binding peaks identified by SPI-seq and DSBCapture-seq. The resultant gene sets from rGREAT were used as the input of the R package fGSEA[40] to conduct cutting edge analyses.

**Mass-spectrometry proteomics with BioID**

Exponentially growing T-47D cells harbouring lentiviral inducible cassettes encoding BirA*-A3B, A3B-BirA* and BirA*-RFP were treated with 0.5 µg/ml doxycycline for 24 hours for acute induction of the the expression of BioID bait proteins, which was followed by treatment of 50 µM biotin. Procedures developed from a previous report were followed[42], except for that Dynabeads™ M-280 streptavidin beads were used instead of the MyOne™ Streptavidin C1 beads reported in the earlier version of the article. The affinity purified protein samples were verified by Western blotting and subjected to tryptic digestion and analysed by quantitative mass spectrometry at the Proteomic Facility at Institute of Cancer Research (external website). Data analysis was performed using Scaffold software[43], followed by filtering out common mass-spectrometry contaminants using the CRAPOME database[44] with SAINT-express score[45]. Results from BirA*-RFP bait protein controls, together with 17 historical BioID control experiments recorded in CRAPOME database were used as input for the analysis.

**Antibodies and siRNA oligonucleotides**

The following antibodies were used for this study:

| *Antibody* | *Clone* | *involved methods* | *Provider* | *Cat. #* |
|---|---|---|---|---|
| Flag tag | M2 | IB, co-IP, ChIP, SPI | Sigma | F1804 |
| HA tag | polyclonal | IB | Proteintech | 51064-2-AP |
| GAPDH | polyclonal | IB | Abcam | ab9485 |
| A3B | EPR18138 | IB, co-IP | Abcam | ab184990 |
| V5 tag | SV5-Pk1 | IB | Abcam | ab27671 |
| DDX5 | polyclonal | IB | Abcam | ab21696 |
| α-tubulin | polyclonal | IB | Abcam | ab4074 |
| DNA:RNA hybrid | S9.6 | ssDRIP, slot blot, co-IP | KeraFAST | ENH001 |
| CSB | E-18 | IB | Santa Cruz | sc-10459 |
| TDRD3 | 2E11 | IB, co-IP | Millipore | MABE1042 |
| TDRD3 | polyclonal | ChIP | Proteintech | 13359-1-AP |
| TOP3B | monoclonal | IB, co-IP | Abcam | ab56445 |
| FMR1 | polyclonal | IB | Proteintech | 13755-1-AP |
| $H3^{R17me2a}$ | polyclonal | ChIP | Active Motif | 39709 |
| CARM1 | polyclonal | ChIP | Active Motif | 39251 |
| Streptavidin-HRP | n/a | IB, slot blot | Pierce | 21130 |

For siRNA, A3B knockdown was performed with siRNA sequence reported by Periyasamy et al.[46], CSB with siRNA reported by Sollier et al.[47], and TDRD3 with siRNA reported by Peng et al.[48].

**qPCR quantification**

qPCR was performed on an Applied Biosystems ViiA 7 thermo cycler using POWER SYBR-Green master mix (Thermo Fisher). The following primer pairs were used in this study:

| Gene | Coordinates (GRCh37) | 5'primer | 3'primer |
|---|---|---|---|
| CISH | chr3:50642686-50643064 | ACCTGGAGGAAGCGTGCCATC | AGCCACGTGCCTTCCCTGTTAC |
| PDPR | chr16:70170509-70170872 | AGAGGTGTGTGCCAAGACATCTGG | TTCAAGACCAGCCTGGCTAACATG |
| PGR | chr11:100904669-100905087 | TTCTGGGACTAGGCCAGCAGTC | AAGCTTGTCCGCAGCCTTATGC |
| RARA | chr17:38478476-38478879 | AGCACAAAAGGCAGGGGAGAAG | AGGCAAGCAAGGTCCCAACTG |

| | | | |
|---|---|---|---|
| AGR3 | chr7:16919998-16920357 | ACCATGTTGGCCAGGCTGATC | AGCACTTTGGGAGGCCGAAGC |
| CABLES1 | chr18:20840360-20840777 | CTGAACAGCTGGCCCCTTGC | AAGCTTGCAGCAGGGCAGAAAG |
| PGR- | chr11:101002083-101002454 | TCAGGACAGCATTGCCAGGTAGTC | ACCTTGTGCCTCAGTTTTCCCAAC |
| RARA- | chr17:38527751-38528130 | ATTGGTCCCCCACGCTGACATG | AGCAGCTAATGGGGGCAAAGAC |

**Datasets used in this study**

The following datasets on public domain was used in this study:

| Dataset name | Reference # | source |
|---|---|---|
| GC-skew (GRCh19) | n/a | R-loop DB |
| T-47D ESR1 ChIP-seq | GSE148277 | GEO |
| T-47D GRO-seq | GSE128452 | GEO |
| T-47D RNAPII ChIP-seq | GSE105793 | GEO |
| T-47D Repli-Seq | ENCFF440QFG | ENCODE |
| T-47D Mnase-seq | GSE74308 | GEO |
